# Proteomic Analysis of Breast Cancer Subtypes Identifies Stromal Protein Profiles that Contribute to Aggressive Malignant Behavior

**DOI:** 10.1101/2025.01.21.634187

**Authors:** Jordan B. Burton, Philippe Gascard, Deng Pan, Joanna Bons, Rosemary Bai, Chira Chen-Tanyolac, Joseph A. Caruso, Christie L. Hunter, Birgit Schilling, Thea D. Tlsty

## Abstract

Breast cancer manifests as multiple subtypes with distinct patient outcomes and treatment strategies. Here, we optimized proteomic analysis of Formalin-Fixed Paraffin-Embedded (FFPE) specimens from patients diagnosed with five breast cancer subtypes, luminal A, luminal B, Her2, triple negative (TNBC) and metaplastic breast cancers (MBC), and from disease-free individuals undergoing reduction mammoplasty (RM). We identified and quantified ∼6,000 protein groups (with >2 peptides per protein) with significant changes in over 26% of proteins comparing each cancer subtype with control RM. Stringent statistical filters allowed us to deeply mine 576 significant conserved protein changes shared by all subtypes and protein changes unique to each subtype. The most aggressive subtype, MBC, revealed exacerbated stromal stress responses, as illustrated by a collagenolytic extracellular matrix (ECM) and immune participation biased towards neutrophils and eosinophils. Immunostaining of breast tissue sections confirmed differences across subtypes, in particular, a strong upregulation of SERPINH1, neutrophil-specific myeloperoxidase and eosinophil cationic protein in MBC. In summary, we present deep proteomic, digitalized protein abundance profiles, generated from FFPE breast cancer tissues, that revealed significant changes in ECM and cellular proteins.

**Statement of Significance of the Study:** This study is significant as it discovered deep proteomic signatures for the highly aggressive and malignant metaplastic breast cancer (MCB) which is now considered a fifth subtype based upon its remarkable intra-tumoral heterogeneity that illustrates its unique cell plasticity. To efficiently analyze formalin-fixed paraffin-embedded (FFPE) breast tissues from patients with different breast cancer subtypes and disease-free individuals, we optimized a novel workflow in which we combined paraffinization and Folch extraction. We identified and confidently quantified ∼6,000 protein groups. We were able to find robust changes in extracellular matrix (ECM) with cancer, even though no ECM enrichments were performed. Interestingly, despite the relatively small human cohort size (42 patients), distinct protein signatures emerged throughout all cancer subtypes – common and unique – with remarkable statistical significance for many cancer-relevant proteins and pathways. This Pilot study indicates the hypothesis that an altered stroma can dictate epithelial tumor cell fate. We also observed that MBC was characterized by an especially immunosuppressed tumor environment. We do note the limitation of the relatively small cohort size of our study, and in the future additional patient cohorts will be needed to further validate our findings.

## 1 | INTRODUCTION

Breast cancer afflicts approximately 1 in 8 women over their lifetime in the United States (1). It is becoming increasingly apparent that improvement of patient outcomes and customization of effective treatment plans for precision medicine will rely on a better understanding of the cellular and intricate molecular changes in tissue ecology that manifest in each breast cancer subtype. Historically, classification of breast cancer subtypes was based upon epithelial traits, i.e. expression (or not) of Estrogen Receptor (ER), Progesterone Receptor (PR) and human epidermal growth factor receptor 2 (Her2). This led to the identification of four subtypes of breast cancers: ER+/PR+/Her2-(Luminal A; LumA), ER+/PR+/Her2+ (Luminal B; LumB), ER-/PR-/Her2+ (Her2) and ER-/PR-/Her2- (Triple Negative Breast Cancer; TNBC) (2).

Although originally considered as part of the TNBC subtype because of its marker profile, Metaplastic Breast Cancer (MBC) is now considered a fifth subtype based upon its remarkable intra-tumoral heterogeneity that illustrates its unique cell plasticity (lineage promiscuity). Indeed, MBCs present with unique regions of metaplasia (not seen in other breast cancer subtypes) that can adopt either spindle, squamous, chondroid or even osseous differentiation (3). For example, MBCs can contain morphologically and immunohistochemically validated sectors of multiple tissue types, such as pancreatic, bone, and brain (Supplemental Fig. S1*A*). Mirroring its histological heterogeneity, MBC is also characterized by genetic heterogeneity that further distinguishes it from TNBC (4). Thus, MBC is highly enriched for *PIK3CA*/*PIK3R1* and *Ras-Map kinase* pathway aberrations compared to TNBC (61% vs.14% and 25% vs 7%, respectively). In addition, MBC typically harbors a high frequency of *TP53* (64%) and *TERT* promoter (25%) mutations. Complementing these findings, a recent study analyzing snap frozen tumor samples documented heterogeneous proteomic profiles for MBC depending on histological type (squamous, spindle or sarcomatoid), when compared to disease-free breast and TNBC (5). Although paraffinized FFPE samples often pose additional challenges and difficulties for proteomic workflows compared to analyses of fresh samples, in our study we were able to analyze the five breast cancer subtypes in great analytical depth.

The epithelial distinctions that typify MBC also extend to stromal distinctions although these are currently understudied in each of the breast cancer subtypes. Notably, MBC frequently expresses immune checkpoint markers indicative of an immunosuppressive microenvironment (6). Moreover, a recent study has established that stromal heterogeneity might explain the increased incidence of MBC in the African American population (7). These recent provocative reports from Kalaw et al., 2020 (6) and Kumar et al., 2023 (7) extend the pioneering work of Finak et al., 2008 (8) who first showed that stromal gene expression could predict clinical outcome in breast cancer. Indeed, molecular profiling of microdissected stroma from primary breast tumors enabled the Park team (8) to identify a stroma-derived prognostic predictor (SDPP) that stratified disease outcome independently of standard clinical prognostic factors and published predictors. The SDPP relies on a panel of genes involved in immune, angiogenic and hypoxic responses, supporting an instructive role of the stroma in tumorigenesis. This instructive function of the stroma on epithelial cell plasticity, i.e., on the epithelial cell propensity to generate metaplastic lesions, has been documented in other metaplastic cancers, such as lung squamous cell carcinoma (9), esophageal adenocarcinoma (10) and gastric adenocarcinoma (11).

In this study, we contrasted proteomic signatures across human breast cancer subtypes, including LumA, LumB, Her2, TNBC and MBC, with disease-free breast tissues, to identify protein signatures common to all cancer subtypes, but also signatures specific to each subtype, with a particular focus on MBC. Technologically, we demonstrate a highly efficient workflow that combined FFPE protein extraction protocols, specifically tailored and adjusted for breast tissues that exhibit high fat content, with quantitative and comprehensive proteomic data-independent acquisitions (12–15). Our goal was to gain biological insights into the contribution of the stroma (including both cellular and acellular components, such as the ECM) to the progression of these breast cancer subtypes. We identified unique (or enriched) signatures associated with the most extreme disruption in tissue homeostasis observed in MBC and validated these signatures by immunohistochemistry. Such candidate markers and signatures may, in the future, lead to potential therapeutic targets.

## 2 | MATERIALS AND METHODS

### 2.1 | Chemicals

Acetonitrile (AH015) and water (AH365) for HPLC were obtained from Burdick & Jackson (Muskegon, MI). Qiagen supplied the Qproteome FFPE Tissue Kit (37623). Reagents for protein chemistry purchased from Sigma Aldrich (St. Louis, MO) include heptane (34873), methanol (34860), ethanol (459844), beta-mercaptoethanol (444203), chloroform (366927), sodium dodecyl sulfate (436143), triethylammonium bicarbonate buffer 1.0M, pH 8.5 (T7408), iodoacetamide (I1149), dithiothreitol (D9779), phosphoric acid (49685), and formic acid (94318-50mL-F). BCA protein assays (23227) and PNGase F (A39245) were obtained from Thermo Fisher Scientific (Waltham, MA). Promega (Madison WI) provided sequencing grade trypsin (V5117). S-trap mini columns were obtained from Protifi (Fairport, NY). Empore C18 resin as 47 mm disks (051115–08807) from 3M (St. Paul, MN) were used for Stage Tips generated in house (3 disk per Stage Tip). Biognosys (Schlieren, Switzerland) supplied the iRT peptides (iRT-Kit).

### 2.2 | Collection of human breast cancer samples

Tissues were obtained from the Cooperative Human Tissue Network Western Division (Nashville, TN) and mid-Atlantic Division (Charlottesville, VA) for FFPE tissue blocks. A panel of formalin fixed paraffin-embedded human breast tissue specimen consisting of 7x ER+/PR+/Her2- (luminal A), 7x ER+/PR+/Her2+ (luminal B), 7x ER-/PR-/Her2+ (Her2+), 7x ER-/PR-/Her2- (triple negative; TN) and 7x metaplastic breast cancers (MBCs) as well as 7x disease-free reduction mammoplasty (RM) specimen used as reference, were processed for proteomics analysis. The age range of patients for these samples was from 22 – 92 years with an average age of 57 years. All obtained human tissue samples were FFPE conserved. Further human cohort details, including sex, race, specimen weight, metastatic outcome, necrosis (%), lesion (%), stroma (%), cellularity (%), acellular (%), stage, grade, and additional comments can be found in Supplemental Table S1. For FFPE Breast Tissue Multiplex Immunohistochemistry (mIHC) see Supplemental Methods.

### 2.2 | FFPE Breast Tissue Deparaffinization and Protein Extraction

Three 20 µm-thick serial sections were cut from a 1 cm^2^ block of FFPE tissue, excess paraffin was cut from the periphery of the tissue area, and each tissue was placed into a 1.5 mL tube. The sections were deparaffinized by adding 0.5 mL of heptane and vortexing vigorously for 10 seconds, followed by incubation at room temperature for 1 hour. Next, 25 µL of methanol were added and the samples were vortexed and centrifuged at 9,000 x g for 2 minutes to pellet the tissue. The supernatant was removed and discarded using a pipet, and the pellet was air-dried for 5 minutes. Then, 100 µL of Extraction Buffer EXB Plus (Qiagen) supplemented with beta-mercaptoethanol were added to the tube containing the pellet and the mixture was vortexed before and while incubation on ice for 5 minutes (16). The sample was subsequently incubated in a Thermomixer at 100 °C for 20 minutes, followed by incubation at 80 °C for 2 hrs with agitation at 750 rpm. Each sample was cooled at 4 °C for 1 minute and centrifuged at 14,000 xg for 15 minutes at 4 °C. The supernatant, which contained extracted proteins, was transferred to a new 1.5 mL tube. The average concentration of proteins in the supernatant was determined to be approximately 2 mg/mL using a BCA protein assay.

Aliquots of 40 µL of the supernatant were taken for additional sample cleanup and digestion (16). The samples were mixed (vortexed) with 200 µL of methanol, and centrifuged at 9000 x g for 10 seconds. Next, 50 µL of chloroform were added to the samples, which were mixed and centrifuged at 9000 x g for 10 seconds to pellet the precipitated proteins. The top layer was removed, and 150 µL of water were added to the samples, which were mixed vigorously and centrifuged at 9000 x g for 1 minute to form an interphase containing protein. This process was repeated with the addition of 150 µL of water, vortexing, and centrifuging at 9000 x g for 1 minute. The top layer was removed and discarded, and 300 µL of methanol were added to the samples, followed by vigorous mixing and centrifuging at 9000 x g for 2 minutes. The supernatant was removed from the protein pellet and discarded, and an additional 300 µL of methanol were added to the samples, which were mixed and centrifuged at 9000 x g for 2 minutes. The supernatant was again removed and discarded, and 1 mL of ethanol was added to each sample, which were mixed and centrifuged at 9000 x g for 2 minutes to pellet the protein. The supernatant was removed and discarded, and the process was repeated with the addition of ethanol. The final pellet was resuspended in 50 µL of 5% sodium dodecyl sulfate in 50 mM triethylammonium bicarbonate (TEAB) through vigorous mixing and incubated at room temperature for 1 hour.

### 2.3 | Protein digestion and desalting

Samples were treated with 20 mM dithiothreitol in 50 mM TEAB, pH 7, at 56 °C for 10 minutes. The samples were then cooled to room temperature and allowed to sit for an additional 10 minutes. Next, the samples were alkylated with 40 mM iodacetamide in 50 mM TEAB (pH 7) at room temperature for 30 minutes in the dark. The samples were then acidified by adding 6.5 µL of 12% phosphoric acid, and a fine precipitate was formed after the addition of 500 µL S-trap buffer and mixing by inversion. The entire mixture was added to S-trap mini spin columns (Protifi) and centrifuged at 4,000 xg for 20 seconds or until fully eluted. The samples were then washed with an additional 400 µL of S-trap buffer. Next, the samples were incubated with sequencing grade trypsin (Promega) dissolved in 50 mM TEAB (pH 7) at a 1:25 enzyme:protein ratio for 1 hour at 47 °C. Additional trypsin was added at the same ratio and the proteins were allowed to digest at 37 °C overnight. The peptides were eluted from the S-trap columns and dried by centrifugal evaporation, and then resuspended in 100 µL of 50 mM TEAB (pH 7). The peptides were then deglycosylated by the addition of 3 µL 1,500 U PNGase F and incubated at 37 °C for 3 hours. The samples were brought to a final concentration of 1% formic acid in water using 10% formic acid and desalted using in-house Stage Tips. The desalted elutions were dried by centrifugal evaporation and resuspended in 20 µL of 0.2% formic acid in water. Finally, indexed Retention Time Standards (iRT, Biognosys) were added to each sample according to the manufacturer’s instructions (17).

### 2.4 | Mass spectrometric analysis

Reverse-phase HPLC-MS/MS data was acquired using a Waters M-Class HPLC (Waters, Massachusetts, MA) connected to a ZenoTOF 7600 system (SCIEX, Redwood City, CA) with an OptiFlow Turbo V Ion Source (SCIEX) equipped with a microelectrode (18). The chromatographic solvent system consisted of 0.1% formic acid in water (solvent A) and 99.9% acetonitrile, 0.1% formic acid in water (solvent B). Digested peptides (4 µL) were loaded onto a Luna Micro C18 trap column (20×0.30 mm, 5 µm particle size; Phenomenex, Torrance, CA) over a period of 2 minutes at a flow rate of 10 µL/min using 100% solvent A. Peptides were eluted onto a Kinetex XB-C18 analytical column (150×0.30 mm, 2.6 µm particle size; Phenomenex) at a flow rate of 5 µL/min using a 120 minute microflow gradient, with each gradient ranging from 5 to 32% solvent B using the following paradigm: digested peptides were loaded at 5% B and separated using a 120 min linear gradient from 5 to 32% B, followed by an increase to 80% B for 1 min, a hold at 80% B for 2 min, a decrease to 5% B for 1 min, and a hold at 5% B for 6 min. The total HPLC acquisition time was 130 minutes. The following MS parameters were used for all acquisitions: ion source gas 1 at 10 psi, ion source gas 2 at 25 psi, curtain gas at 30 psi, CAD gas at 7 psi, source temperature at 200°C, column temperature at 30°C, polarity set to positive, and spray voltage at 5000 V.

All human samples were acquired in data-independent acquisition mode (DIA) analysis with two technical replicates (2 injections) for each biological sample replicate as described in detail in the online methods. Briefly, the DIA-MS method on the ZenoTOF 7600 system is comprised of A survey MS1 scan (mass range: 395-1005 m/z), with an accumulation time of 100 ms, a declustering potential of 80 V, and a collision energy of 10 V. MS2 scans were acquired using 80 variable width windows across the precursor ion mass range (399.5-1000.5 m/z), with an MS2 accumulation time of 25 ms, dynamic collision energy enabled, charge state 2 selected, and Zeno pulsing enabled (total cycle time 2.5 seconds, Supplemental Table S6).

### 2.5 | DIA data processing with Spectronaut

All data files were processed with Spectronaut v16 (version 16.0.220524.5300; Biognosys) performing a spectral library using the *Homo sapiens* pan-human spectral library with 20,526 entries, accessed on 08/28/2015 (19). Dynamic data extraction parameters and precision iRT calibration with local non-linear regression were used. Trypsin/P was specified as the digestion enzyme, allowing for specific cleavages and up to two missed cleavages. Methionine oxidation (+15.995 Da), protein N-terminus acetylation (+42.011 Da), and deamidation of asparagine (+0.984 Da) were set as dynamic modifications, while carbamidomethylation of cysteine (+57.021 Da) was set as a static modification. Protein group identification (grouping for protein isomers) required at least 2 unique peptides and was performed using a 1% false discovery rate (FDR) estimated with PyProphet for peptide spectrum match, peptide and protein level. Protein quantification was based on the peak areas of extracted ion chromatograms (XICs) of 3 – 6 MS2 fragment ions, specifically b- and y-ions, with automatic normalization and 1% q-value data filtering applied. Relative protein abundance changes were compared using the Storey method with paired t-tests and p-values corrected for multiple testing using group wise testing corrections (20). The quantification significance level was as follows: q-value less than 0.001 and absolute Log2(fold-change) greater than 0.58 when comparing breast cancer types to disease-free RM specimen. Details for additional data processing and visualizations as well as additional statistical details are provided in the Supplemental Methods.

## 3 | RESULTS

### 3.1 | Proteomic Workflow and Optimization of Tissue Preparation

Our study utilized a comprehensive proteomic quantification, profiling and digitalization approach: data-independent acquisition (DIA) to analyze formalin-fixed paraffin-embedded (FFPE) tissue specimens representative of five distinct breast cancer subtypes (Fig. 1, *A* and *B*) which include Luminal A, Luminal B, HER-2-expressing, Triple Negative and metaplastic breast cancer (LumA, LumB, Her2+, TNBC and MBC, respectively) (12, 13, 21). Archived FFPE tissue samples are an invaluable source of clinical research material that often cannot otherwise be obtained. However, the extraction of soluble proteins from FFPE tissues remains challenging because the fixation process can result in the cross-linking of proteins, often leading to low yields, protein degradation, and reduced accessibility of membrane-bound and low-abundant proteins (22). We optimized our extraction procedures to improve mass spectrometric acquisition as described in Materials and Methods. Briefly, FFPE specimens were deparaffinized, followed by Folch-extraction (chloroform/methanol/water extraction) prior to subsequent proteolytic digestion using trypsin and de-glycosylation with PNGase F (Fig. 1*C*) (16). It should be noted that for our experimental paradigm previously reported methods for FFPE deparaffinization were not efficient enough for our specific breast tissue samples to provide fully delipidated samples for optimal MS analysis. Specifically, initial sample preparation methods only using deparaffinization failed with inefficient protein extraction (identifying only 50-100 proteins per sample and exhibiting poor chromatography and peak shapes which was not appropriate for any type of quantification). The initially inefficient workflow – we hypothesize - may also have been resulting from challenges related to breast tissue composition containing a lot of adipose tissue. However, the optimization and addition of multiple Folch extraction steps (after deparaffinization) improved sample quality by successfully removing any residual lipid content associated with the fatty and adipocyte-rich nature of the breast tissue. As a result of the above tailored protocol to extract proteins from FFPE embedded human breast tissue we were now able to identify and quantify 5,858 protein groups (with each protein identified with at least 2 unique peptides). The 5,858 identified protein groups were localized to multiple organelles, as illustrated by the top 15 cellular compartments in Supplemental Fig. S2. Indeed, the analysis of whole tissue from FFPE specimens allowed isolation of proteins that were not simply restricted to the ECM and cytoskeleton (such as when enriching for ECM as we previously have reported for lung cancer (14)), but that also extended to structural proteins (cytoplasmic, ER, Golgi apparatus and nuclear-associated proteins) as well as functional proteins (such as enzymes) typically found intracellularly.

**FIG. 1.**
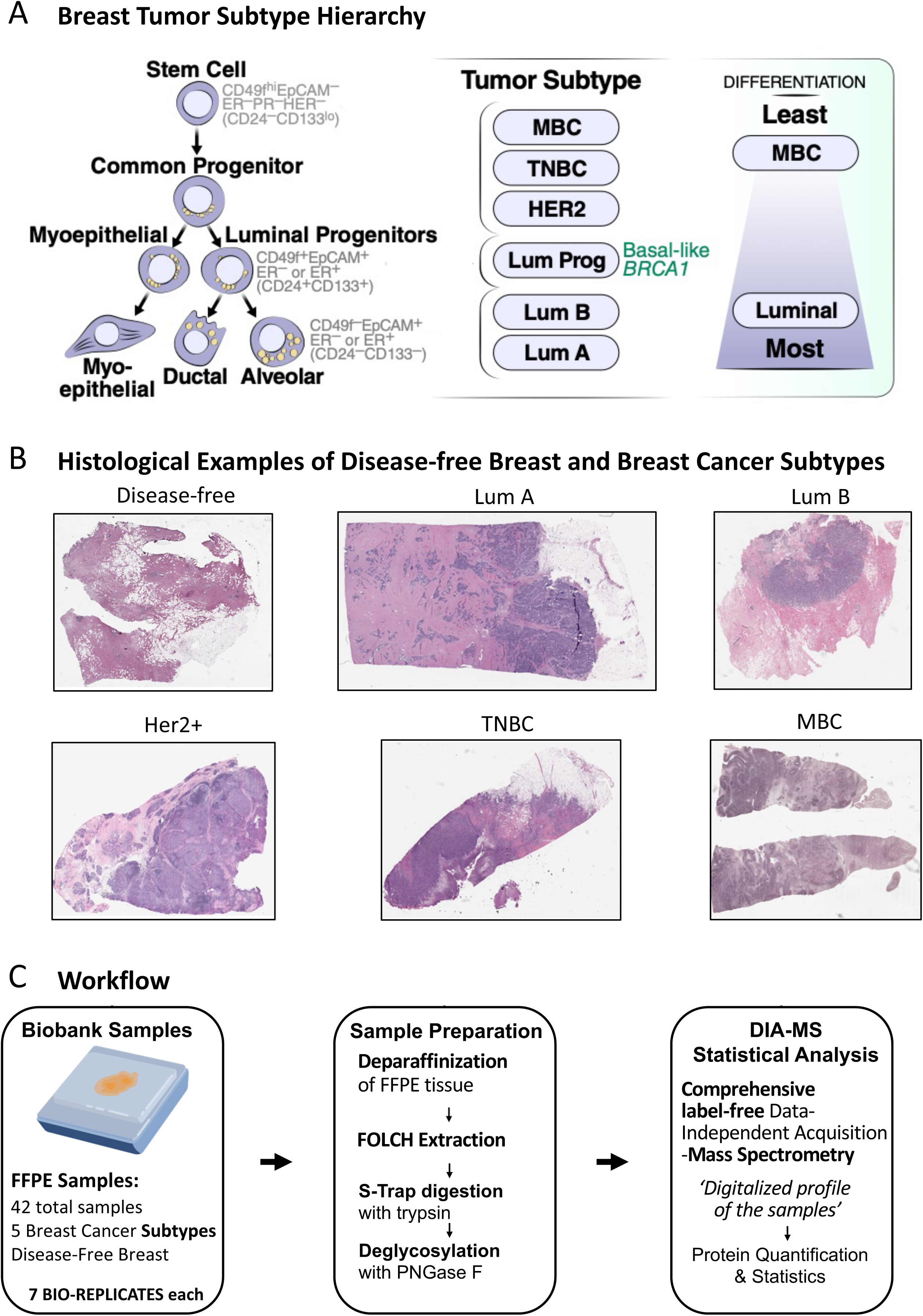
Breast cancer subtypes and workflow for proteomic analysis of human breast tissue FFPE specimens. *A*, Breast cancer subtypes exhibit a continuum from well differentiated to poorly differentiated phenotype, *adapted from Lim et. al. Nat. Med. 2009* (*2*). The 5-year survival rates are 55% for metaplastic breast cancer (MBC), 77% for triple negative breast cancer (TNBC), 83% for HER2 positive breast cancer (HER2), 92% for luminal B breast cancer (LumB), and 89% for luminal A breast cancer (LumA) (1). *B*, Showcase of one hematoxylin and eosin stained FFPE tissue for representative sample subtypes, including a reduction mammoplasty (RM) control. *C*, A workflow for processing and acquiring proteomic data from human breast FFPE tissue is shown. Briefly, 42 FFPE samples were obtained with known pathologies. Tissue scrolls were collected and deparaffinized before protein extraction. Folch extracted proteins were digested with trypsin and deglycosylated with PNGase F.

A panel of FFPE human breast tissue specimens (Supplemental Table S1) consisted of 7 LumA, 7 LumB, 7 Her2+, 7 TNBC and 7 MBC diagnosed patients, as well as 7 disease-free reduction mammoplasty control specimens, and was processed for proteomic analysis (Supplemental Fig. S1, *B-G*). All samples were acquired by data-independent acquisitions (DIA) with efficient quantification statistics when comparing the different conditions (Fig. 1*C*) (12, 13, 21). The identification and quantification of a total of 5,858 protein groups (with at least 2 peptides per protein) is shown in Supplemental Table S2-*A* and abided to stringent proteomic guidelines (23). Significantly altered protein groups were determined and filtered with an absolute log2-fold change ≥0.58 and a very stringent q-value ≤0.001 (quantification confidence) when comparing each breast cancer subtype specimen with disease-free breast specimens, respectively. Comparisons of each of the 5 breast cancer subtype with the disease-free breast tissues are displayed in volcano plots indicating a very large percentage (∼25%) of significantly changed proteins (Fig. 2, *A-E* and Supplemental Table S2, *B-F*). We further mined the data sets and investigated similarities (Supplemental Fig. S3) and unique differences observed in each of the five breast cancer subtypes (Fig. 2*F*).

**FIG. 2.**
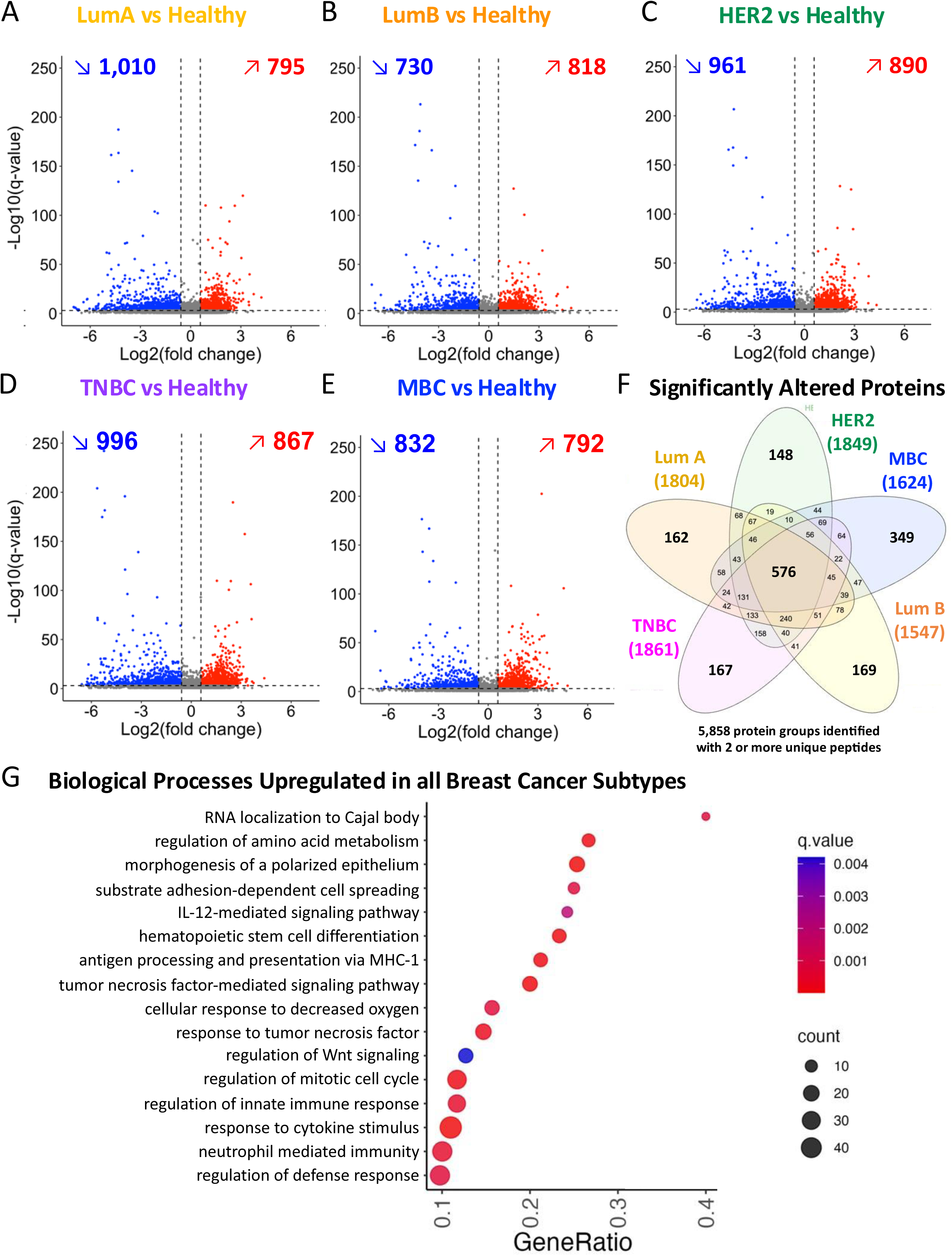
Breast Cancer Subtype Proteomes revealed tissue remodeling. Volcano plots of the 5,858 protein groups identified with 2 or more unique peptides show proteome remodeling for comparisons to disease-free tissue. Significantly changing proteins are indicated in red (up-regulated) or blue (down-regulated): significantly altered proteins were selected with strict q-values ≤ 0.001 and absolute fold changes at |Log2(FC)| ≥ 0.58; non-significantly changing proteins are indicated in grey. *A*, Luminal A (ER+PR+Her2-) breast cancers showed 1,805 proteins with significantly altered levels: with 1,010 downregulated proteins (blue), and 795 upregulated proteins (red). Of these significantly changing proteins 244 proteins (13.5%) only changed significantly in the luminal A subtype, but not in the other subtypes. *B*, Luminal B (ER+PR+Her2+) breast cancers showed 1,548 significantly altered proteins with 730 downregulated proteins (blue), and 818 upregulated proteins (red). Of these significantly changing proteins 169 proteins (10.9%) only changed significantly in the luminal B subtype, but not in the other subtypes. *C*, Her2+ (ER-PR-Her2+) breast cancers showed 1,851 significantly altered proteins (including 148 unique proteins; 8%) with 961 downregulated proteins (blue), and 890 upregulated proteins (red). Of these significantly changing proteins 148 proteins (8.0%) only changed significantly in the Her2+ subtype, but not in the other subtypes. *D*, TNBCs (ER-PR-Her2-) showed 1,861 significantly altered proteins with 996 downregulated proteins (blue), and 867 upregulated proteins (red). Of these significantly changing proteins 166 proteins (8.9%) only changed significantly in the TNBC subtype, but not in the other subtypes. *E*, MBCs showed 1,624 significantly altered proteins (including 349 unique proteins; 21.5%) with 832 downregulated proteins (blue), and 792 upregulated proteins (red). Of these significantly changing proteins 349 proteins (21.5%) only changed significantly in the MBC subtype, but not in the other subtypes. *F*, The Venn diagram showed the overlap of 576 significantly altered protein groups that changed in comparisons of all subtypes vs. control, and significantly altered protein groups that are unique to each cancer subtype (breast cancer subtype vs. disease-free). *G*, Dot plot of the upregulated KEGG biological processes with a q-value ≤ 0.01 and term level ≥ 5 in all breast cancer subtype comparisons with disease-free tissue.

### 3.2 | Conserved Protein Profile Changes Across All Breast Cancer Subtypes

The label-free DIA-MS approach provided proteome quantification and profiling, that revealed 576 significantly altered protein groups with conserved signatures across all breast cancer subtypes (Fig. 2*F* and Supplemental Fig. S3*A*). These shared significant protein changes between all subtypes were related to tumor growth, such as increased transcription and protein translation, changes in metabolic pathways reflecting shifts in lipid and sugar metabolism and mitochondrial function, stress responses, DNA repair and lastly and significantly, large-scale ECM remodeling (Supplemental Table S3-A, ECM signatures are described in more detail below).

### 3.3 | Cell Intrinsic Processes: Metabolism, Morphology, Proliferation, DNA Repair

In general, biological processes that were upregulated in all breast cancer subtypes (Fig. 2*G*) reflected increased energetic and biosynthetic demands for rapid cancer cell proliferation. For example, the upregulation of lactate dehydrogenase A (LDHA), a key checkpoint of anaerobic glycolysis, in the context of a universal stress response, such as the upregulation of heat shock proteins HSP90 and HSP47/SERPINH1, typified such biological processes known to propagate pro-tumorigenic signals in all breast cancers (24, 25). These phenotypes are examples of proteomic results that reflect the biology intrinsic to epithelial cells.

### 3.4 | Widespread Changes Observed Across All Breast Cancer Subtypes

Many of the protein changes observed in our analysis were not confined to epithelial cells but rather extended to stromal components. For example, downregulation of alcohol dehydrogenase 1B (ADH1B) and decorin (DCN) across most breast cancer subtypes (Supplemental Table S3-A) also documented additional significant changes in stromal cell landscape, in this instance fibroblasts. Indeed, the loss of ADH1B has been recently associated with acquisition of a carcinoma associated fibroblast (CAF) state contributing to an interleukin 6 tumor promoting phenotype in colon cancer (26). Loss of stromal CD36, a cell surface scavenger receptor, was observed in all cancer subtypes (Supplemental Table S3-A) and typified the acquisition of a pro-tumorigenic tissue microenvironment that extends to all stromal components through a multicellular coordinated program (27, 28). In addition to altering fibroblast production of collagens and macrophage changes in efferocytosis, the loss of CD36 was recently demonstrated to disrupt vascularization (29), as well as alter lipid metabolism and associated changes in adipose tissue inflammation (30). This measurement validated our previous findings reporting a downregulation of CD36 across human breast cancer subtypes (27) and more recently in lung squamous cell carcinoma (LSCC)(14) and esophageal adenocarcinoma (EAC) (10).

### 3.5 | Extracellular Matrix Remodeling and Signaling

Importantly, we observed shared changes in proteins that suggest alterations in ECM composition and remodeling, which are extrinsic to epithelial cells. ECM proteins are part of a highly dynamic tissue environment. Through control of tissue stiffness, viscoelasticity and degradability, ECM proteins play a pivotal role in regulation of cell behaviors (31). Alterations in ECM have been implicated as key drivers of tumorigenesis and metastasis in all cancers. Examples include breast cancer and chronic inflammation-associated cancers of the lung, i.e. LSCC (14, 32, 33). Consistent with these observations, our proteomic analysis revealed extensive remodeling of the ECM content in the breast tumor microenvironment of all subtypes. Supplemental Fig. S4*A* highlights the 2,023 protein groups quantified with the gene ontology term “Extracellular”, of which 263 proteins are also annotated in the Human Matrisome Database (MatrisomeDB (34), Supplemental Fig. S4*B* and Supplemental Table S4). Of these 263 proteins, 42 proteins were shared in their altered profile amongst the 5 breast cancer subtypes (Fig. S4*C*). The widespread significant downregulation of small leucine-rich, desmosome, and basement membrane proteins (Fig. 3, *A-C*) (35), such as perlecan (HSPG2), desmoplakin (DSP), lumican (LUM), and a multitude of collagens (COLs), implied detrimental degradation of normal ECM composition in all breast cancers. Concurrently, upregulation of factors known to increase epithelial motility and invasion, such as vitronectin (VTN), versican (VCAN), transforming growth factor beta induced (TGFBI), and cathepsin B (CTSB), are postulated to further contribute to loss of tissue integrity and increased tumor cell dissemination. Notably, of all collagens (COL), COL12A1, a structural protein previously implicated in breast cancer metastasis through spatial mass spectrometry imaging (36), was the only collagen to be upregulated in all breast cancer subtypes in our dataset. The upregulation of cross-linking enzymes of the lysyl oxidase (LOX) family, as well as the increase in selected matrix metalloproteases (MMPs), MMP8, MMP9, MMP14, also reflected an extensive tissue microenvironment remodeling favoring tumor cell invasion. Interestingly, it has been recently reported that cleaved collagen I activates a mitochondrial biogenesis pathway, i.e. the discoidin domain receptor 1 (DDR1)-NF-κB-p62-NRF2 signaling axis, that promotes the growth of pancreatic ductal adenocarcinoma (PDAC), while in contrast, intact collagen I triggers the degradation of DDR1 and limits the growth of PDAC, a highly desmoplastic, aggressive and metastatic cancer (37). Thus, increased degradation of collagens and MMP activation, defined as a collagenolytic stroma, may therefore contribute to greater aggressivity of some breast cancer subtypes. While MMP level changes were shared amongst the breast cancer subtypes, the observed changes were not equal and the MMP profile observed in MBC was the one most consistent with a highly aggressive clinical phenotype often observed in chronic inflammation-associated cancers (CIACs) (Supplemental Fig. S5*A*).

**FIG. 3.**
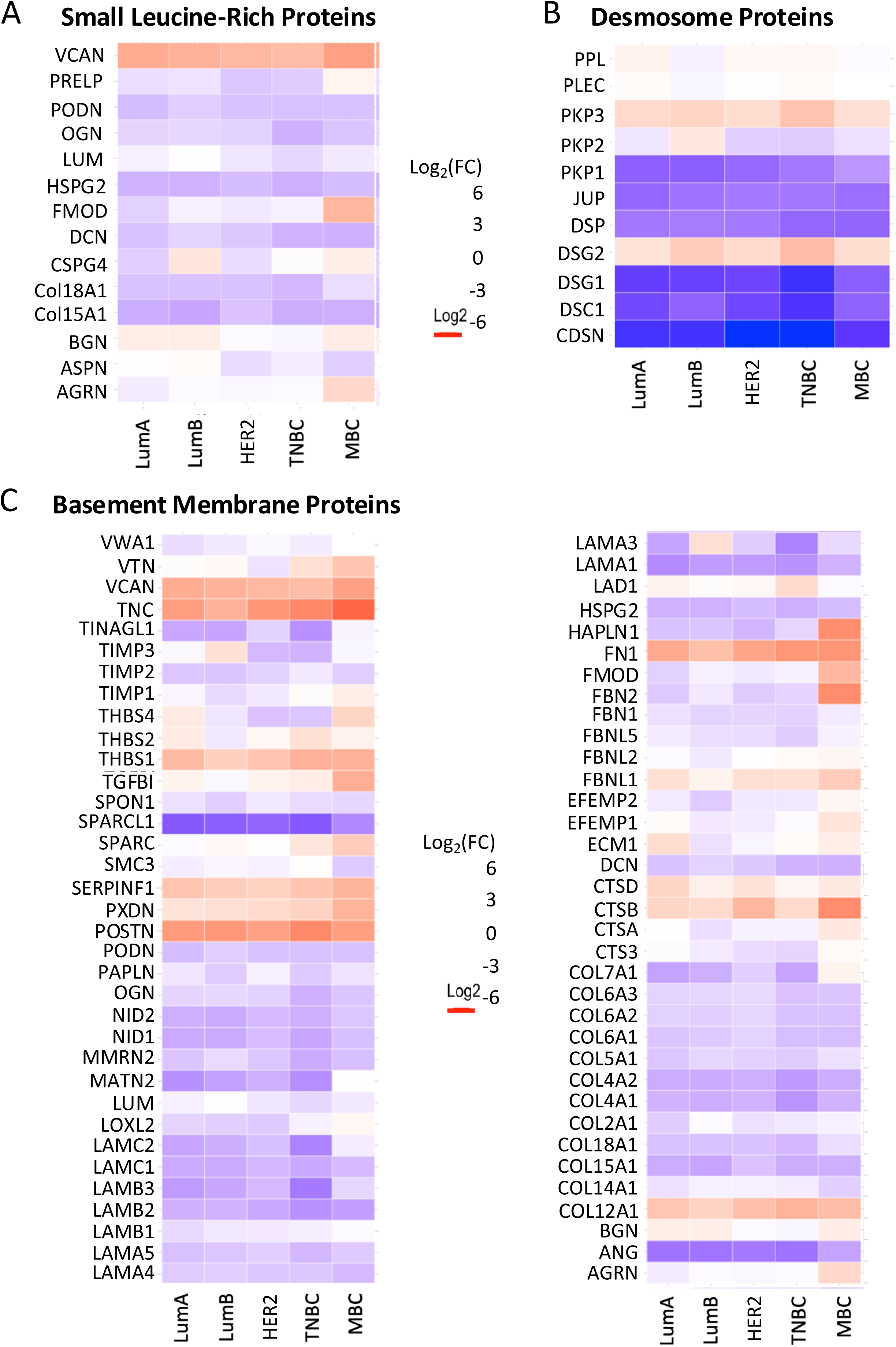
Breast cancers display alterations in tissue microenvironment characterized by quantitative and qualitative changes in ECM proteins. *A*, Heatmap of the significantly altered small leucine-rich proteins. *B*, Heatmap of the significantly altered desmosome proteins. *C*, Heatmap of significantly altered basement membrane proteins. The fold-change scale is displayed for all heatmaps where the positive log_2_(FC) values are shown in red and the negative values in blue.

### 3.6 | Stress Response and Senescence-Associated Secretory Phenotype (SASP)

Injured and diseased tissues often display stress responses characterized by a disruption of homeostasis and a senescence-associated secretory phenotype (SASP). In this context, we detected an upregulation of proteins involved in stress response and influencing ECM processing (Fig. 4). Thus, SERPINH1 (HSP47), a collagen chaperone previously identified and validated by our team as upregulated in LSCC and EAC (14), displayed an increase in levels across all breast cancer subtypes when compared to control tissues. The strong upregulation of SERPINH1 in all breast cancer subtypes, observed to the greatest extent in MBC, was confirmed by immunohistochemical analysis of an extended cohort of specimens including those subjected to proteomic analysis (Fig. 5*A*), while the disease-free tissue showed almost no IHF signal at all. Furthermore, co-staining with the mesenchymal marker vimentin (VIM) confirmed that cells expressing high levels of SERPINH1 were stromal components. Quantitative analysis of VIM+SERPINH1+ cells confirmed a higher density of such cells in MBC and to a lesser extent in TNBC compared to other breast cancer subtypes and disease-free breast (Fig. 5*B* and Supplemental Table S5). Endogenous ‘danger’ signal molecules released by damaged and dying cells, described as damage-associated molecular patterns (DAMPs), activate the innate immune system. Our dataset identified a large set of DAMPs upregulated across breast cancer subtypes including VCAN, TNC, HSPA1A, HSPA5, HSPA8, HSP90AA1, HSP90AB1, HSP90B1 (Fig. 4, *B* and *C*). In contrast with the above-described proteins, which exhibit an upregulation when the cell experiences stress, we also observed a consistent downregulation of some proteins under the same conditions. For example, expression of CD36 (discussed above), was repressed across all breast cancer subtypes, with the most significant decrease noted in MBC tissues when compared to disease-free control (Supplemental Table S3).

**FIG. 4.**
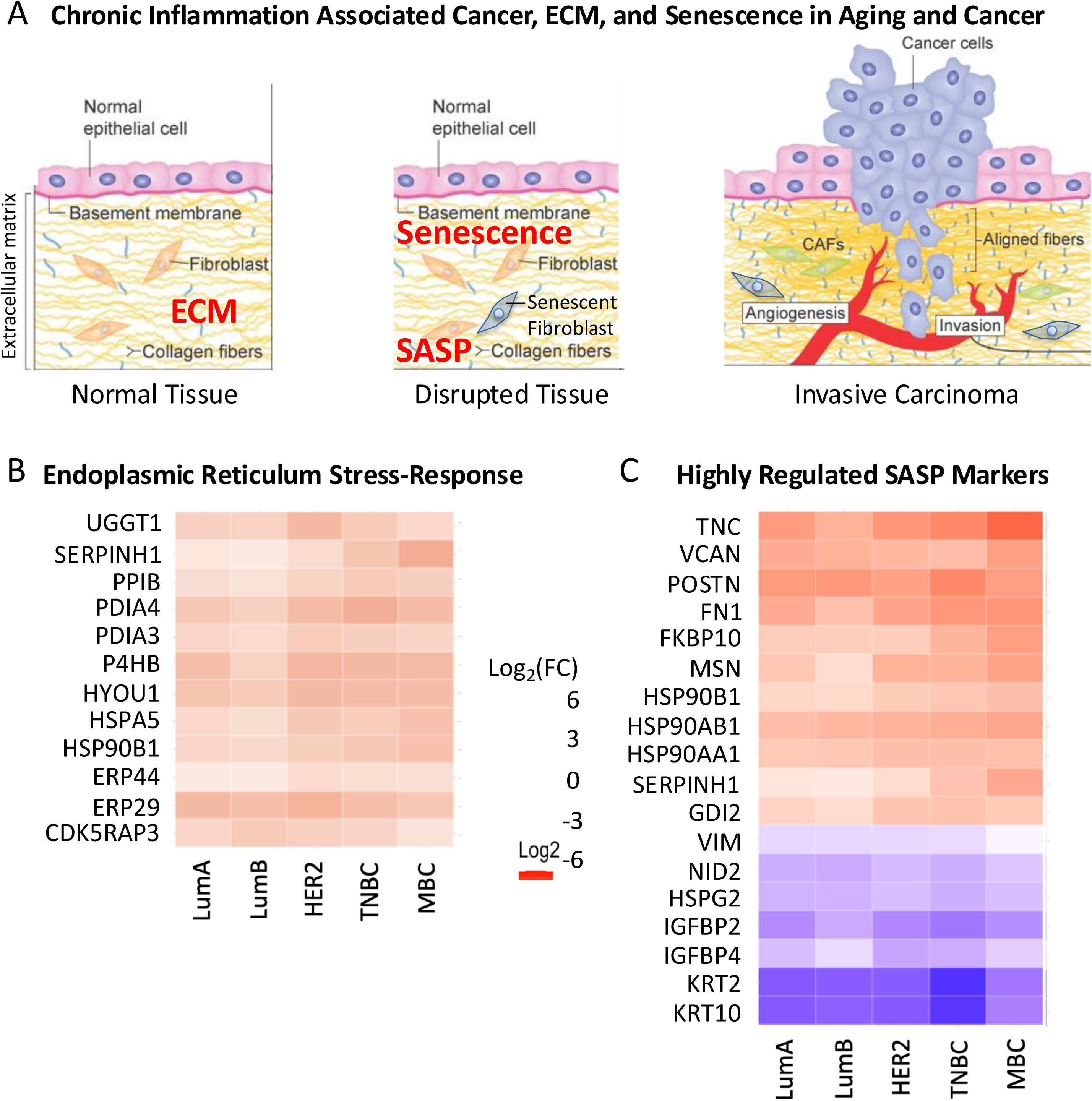
Breast Cancer Subtypes Show an Upregulation of SASP Markers Compared to Disease-Free Samples. *A*, biological overview of the transition from normal tissue to disrupted tissue with increased markers of stress and cellular senescence due to acute inflammation to invasive carcinoma as acute inflammation becomes chronic inflammation. *Adapted from Luthold et al. Cancers, 2022* (110). *B*, Heatmap showing upregulation of stress-response markers from the endoplasmic reticulum. *C*, Heatmap showing higher expression of selected SASP markers in breast cancer subtypes with the worse 5-year survival. Note that all SASP markers are upregulated in the secretome of irradiated IMR 90 cells. Proteins shown have a q-value ≤ 0.001, positive fold change values are displayed in red and negative fold change values are displayed in blue.

**FIG. 5.**
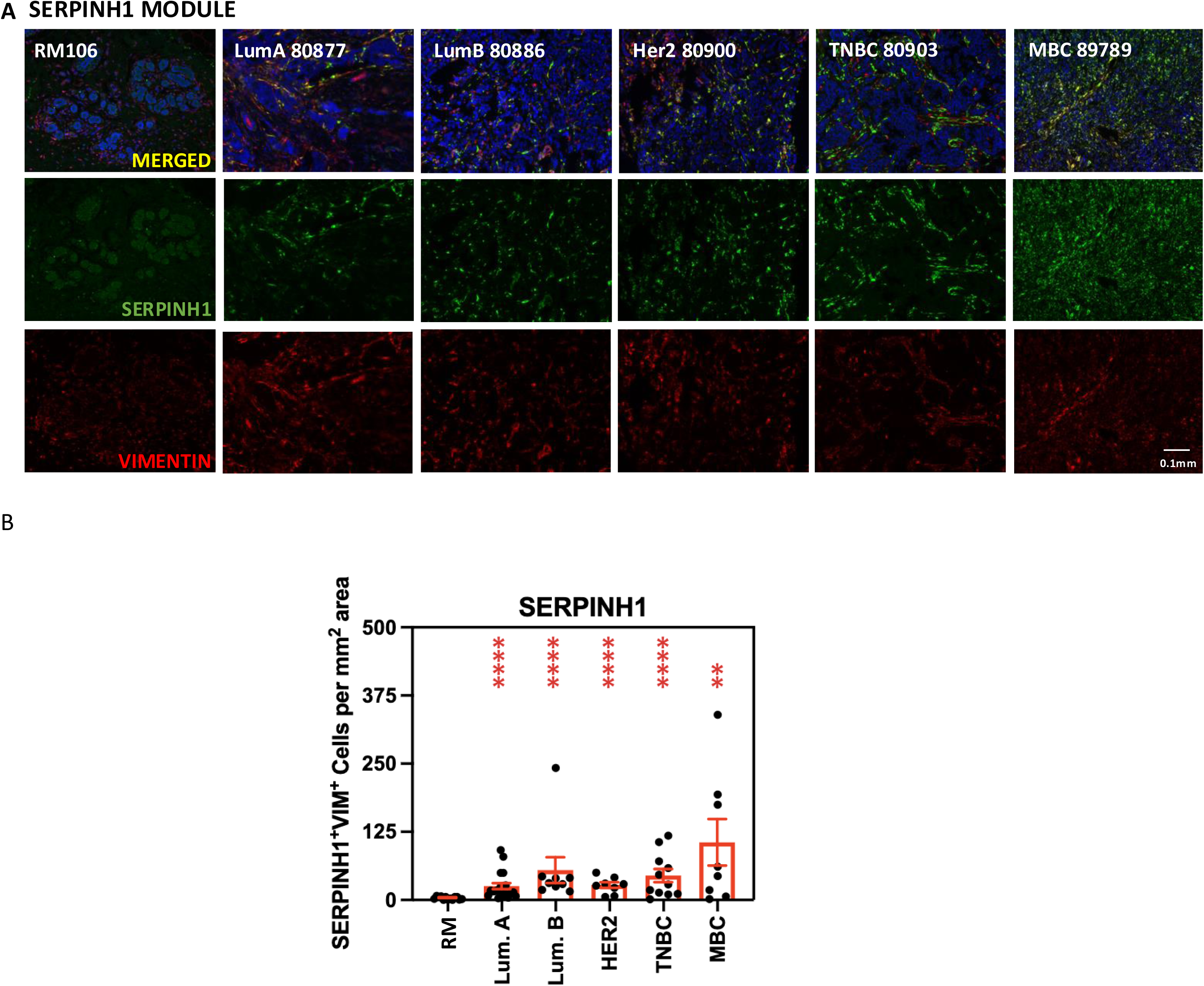
Stromal signature characterized by upregulation of SERPINH1 is exemplified in the least differentiated breast cancers. *A*, Tissue sections from disease-free reduction mammoplasties (RMs) and five breast cancer subtypes (Luminal A (LumA), Luminal B (LumB), Her2-positive (Her2), triple negative (TNBC) and metaplastic (MBC) were sequentially immunostained as described in Methods for collagen chaperone *SERPINH1* detected with FITC and the stromal marker *vimentin* detected with Cy3. Stained slides were counterstained with DAPI and mounted prior to imaging. Representative images (at 20 x magnification) for one specimen of each of the 6 tissue specimen groups described above are shown. Scale bar: 0.1mm. *B*, Quantitative analysis of chaperone SERPINH1. Images of immunostained breast tissue sections were subjected to quantitative analysis (Table S5) as described in Methods. Numbers of SERPINH1+VIM+ cells (fibroblasts) per mm2 of analyzed tissue specimen are displayed for disease-free breast (RM; n=13) and each breast cancer subtype (LumA; n=20, LumB; n=9, Her2; n=8, TNBC; n=11 and MBC; n=7). Degree of significance for each pair comparison (**, ****) are displayed at the top of the panel. P-values: LumA vs. RM: <0.0001; LumB vs RM: <0.0001; Her2 vs. RM: <0.0001; TNBC vs. RM: <0.0001; MBC vs. RM: p=0.0018. Parameters used to generate the graph are shown in Supplemental Table S5.

Both epithelial and stromal cells can senesce prematurely in a pathological context and secrete high levels of proteases, inflammatory cytokines, immune modulators and growth factors (38). Many proteins that were upregulated and identified as stress response proteins, for example SERPINF1, overlap with those described to be secreted by senescent cells (see SASP Atlas (38)). A distinct set of SASP proteins that was significantly altered in the breast cancer subtypes is provided in Fig. 4*C*. Together, these shared alterations fundamentally change the biochemical and biomechanical properties of the matrix and tissue, generating a tumor-supportive landscape tailored for malignant progression. Our findings underscore the vital role of abnormal signaling and ECM remodeling in driving cancer pathogenesis and highlight common tumor adaptations that support malignancy regardless of tissue origin or cancer subtype.

### 3.7 | Identification of Protein Profile Changes Unique to Breast Cancer Subtypes

We observed that the human breast cancer subtypes each showed specific and unique protein marker signatures assessing the significantly altered proteins unique to each comparison of breast cancer subtype to disease-free tissue (Fig. 2*F*, Supplemental Fig. S3*B*. and Supplemental Table S3, *B-F*). As a first step to gain insight into breast cancer subtype-specific signatures, we conducted a Weighted Gene Co-expression Network Analysis (WGCNA) to study biological networks based on pairwise correlations between variables. The WGCNA analysis was implemented to identify modules of highly correlated proteins, potentially sharing common biological functions or regulatory mechanisms (39). These modules were then associated with different breast cancer subtypes, elucidating biological processes specific to each subtype (Supplemental Fig. S6). The modules that showed strong correlation to one or more breast cancer subtypes were selected for further investigation (Fig. 6*A*) as described below. Differentially abundant proteins in each WGCNA module are depicted in Fig. 6, *B-D*.

**FIG. 6.**
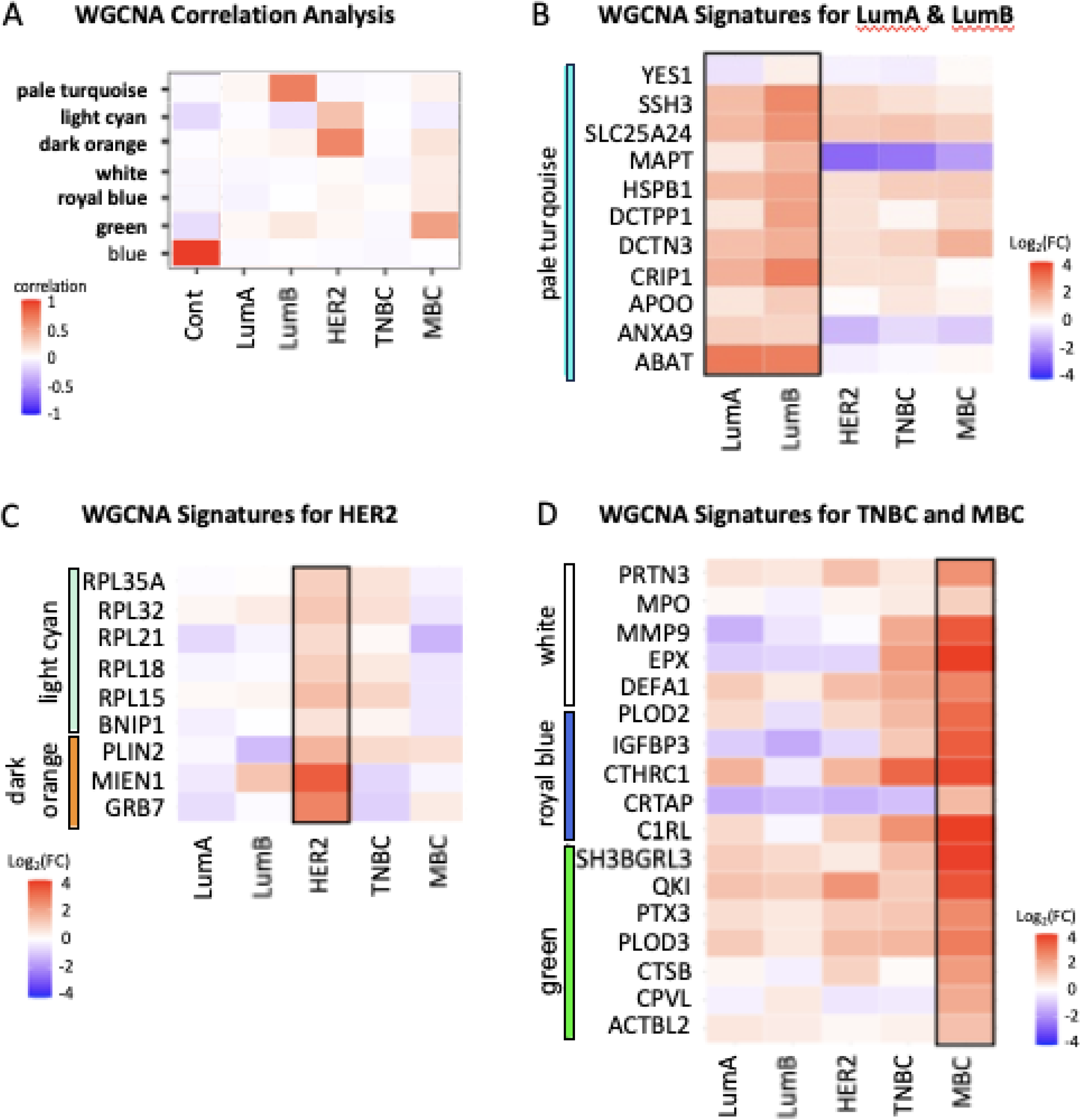
Protein modules of interest identified by Weighted Gene Correlation Network Analysis (WGCNA) define breast cancer subtype-specific proteomic signatures. *A*, WGCNA correlation analysis of select protein modules to breast cancer subtype is displayed. Heatmap of selected significantly altered protein groups from modules defined in WGCNA that are significantly different for *B*, Luminal A and B; *C*, HER2-positive; and *D*, TNBC and Metaplastic breast cancers are shown. The pale turquoise WGCNA module proteins are upregulated signatures for LumA and LumB. The light cyan and dark orange WGCNA module proteins are upregulated in HER2. The WGCNA modules white, royal blue, and green proteins are upregulated in MBC.

### 3.8 | WGCNA Analysis of Luminal A and Luminal B Breast Cancers

The ’pale turquoise’ WGCNA module, specifically associated with LumA and LumB breast cancer subtypes (Fig. 6*B*), featured upregulated proteins implicated in diverse cellular processes. YES proto-oncogene 1, src family tyrosine kinase (YES1), participates in the regulation of critical cellular processes like proliferation, migration, and survival (40). Proteins such as slingshot protein phosphatase 3 (SSH3) and dCTP pyrophosphatase 1 (DCTPP1) are integral to the regulation of the cell cycle and promote cancer cell proliferation (41, 42). Solute carrier family 25 member 24 (SLC25A24) plays a role in cellular metabolism, i.e. mitochondrial function (43), and molecular transport. Heat shock protein family B (Small) member 1 (HSPB1) is correlated with the presence of estrogen and progesterone receptors, and plays a role in Epithelial-Mesenchymal Transition (EMT), cellular stress responses and may be involved in breast cancer metastasis (44). Dynactin subunit 3 (DCTN3) directs cellular trafficking and when upregulated plays a role in breast cancer progression (45). Cysteine rich protein 1 (CRIP1) is involved in cellular growth processes and plays a role in homologous recombination repair, increasing resistance to chemotherapy (46). Apolipoprotein O (APOO) is critical for maintaining lipid transport, and could be involved in breast cancer development (47, 48).

Three proteins in the LumA/LumB module exhibited the significant changes in protein levels between Luminal A and B and the other subtypes. Microtubule associated protein tau (MAPT), a protein that is critical for maintaining cellular structure, has been associated with better patient survival in breast cancer, and was upregulated in LumA and LumB breast cancer subtypes which typically feature a better prognosis for patients and actively downregulated in those with poor prognosis (49). A significant difference was observed in MAPT abundance when compared to more aggressive cancers. Particularly interesting, annexin A9 (ANXA9), was also upregulated only in LumB and LumA subtypes. ANXA9 has been associated with cell signaling, homeostasis, and protects against the immunosuppressive tumor microenvironment, improving patient prognosis (50). The protein coding for 4-aminobutyrate aminotransferase (ABAT), a positive marker for estrogen-receptor α-positive inflammatory breast cancer, is crucial for its role in cellular metabolism and molecular transport (51, 52).

### 3.9 | WGCNA Analysis of HER2+ Breast Cancer

Two WGCNA modules, ’light cyan’ and ’dark orange’, (Fig. 6*C*) were specific to HER2-positive breast cancer. The ’light cyan’ module featured ribosomal proteins (RPLs), essential components in protein synthesis, a process often amplified in rapidly dividing cancer cells (53). Two proteins, which were part of the ‘dark orange’ module, were the most upregulated proteins solely and specifically in the HER2 breast cancer subtype. Notably, the genes encoding these two proteins, migration and invasion enhancer 1 (MIEN1) and growth factor receptor-bound protein 7 (GRB7), are immediately adjacent to the ERBB2 gene that encodes HER2. MIEN1 underscores the aggressive nature of HER2 positive cancers due to its involvement in cell migration and invasion (54). The upregulation of GRB7 highlights the importance of dysregulated signaling in cancer progression due to its role in cell signaling pathways controlling cell growth and survival (55). The ’dark orange’ module also included proteins such as perilipin-2 (PLIN2), associated with lipid metabolism indicative of the altered metabolic state known as the Warburg effect often seen in cancer cells (56).

### 3.10 | WGCNA Analysis of Triple Negative and Metaplastic Breast Cancer

The ’white’ WGCNA module (Fig. 6*D*), primarily consisted of proteins related to neutrophil activity, and drew particular attention due to several significant features. Neutrophil activity was reported in a prior proteomic study in fresh frozen tissues comparing MBC sub-classes to TNBC (5), where it was indicated as a characteristic MBC signature linked with immune-mediated ECM remodeling. Our FFPE tissue dataset expanded on the previous work by demonstrating that such an extensive neutrophil recruitment to the tumor site was a prominent characteristic of MBC, as it was not only less prominent in TNBC, but also not detected in the other subtypes of breast cancer (Fig. 6*D*). The differences in neutrophil recruitment across breast cancer subtypes was validated by immunohistochemistry as discussed below. Additional proteins identified in this ‘white module’, such as proteinase 3 (PRTN3), defensin alpha 1 (DEFA1), and MMP9, have been shown to play a role in neutrophil plasticity (57). These findings suggest a ’fluid cell state’ where not only the epithelial cells, but also stromal cells, such as neutrophils, exhibit plasticity (58). Notably, this remarkable upregulation of neutrophil-specific myeloperoxidase (MPO) and eosinophil-specific eosinophil cation protein (RNAse3) in MBCs, observed to a much lower extent in other breast cancer subtypes, was confirmed by immunohistochemical analysis of the specimens subjected to proteomic analysis (Fig. 7, *A* and *B*). Furthermore, co-staining with the myeloid marker integrin alpha M (ITGAM), a protein also identified in our proteomic analysis as specifically upregulated in MBC, confirmed that most of the cells expressing high levels of MPO or RNAse3 were indeed neutrophils and eosinophils. Quantitative analysis of MPO+ITGAM+ cells and RNAse3+ITGAM+ cells confirmed a higher density of such cells in MBC and to a lesser extent in TNBC compared to other breast cancer subtypes and disease-free breast (Fig. 7*C* and Supplemental Table S5).

**FIG. 7.**
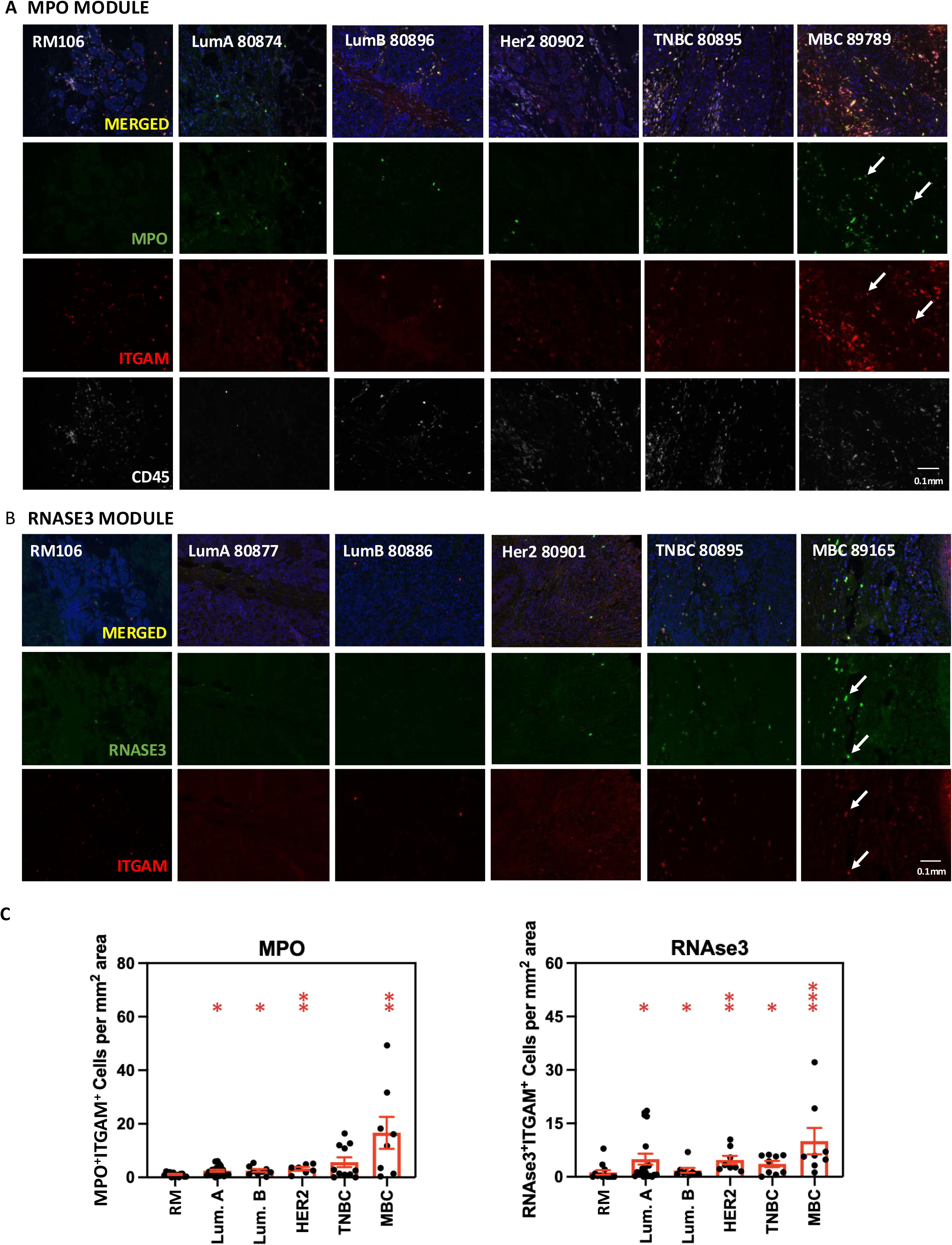
Stromal signatures characterized by upregulation of Myeloperoxidase and RNAse3 are exemplified in the least differentiated breast cancers. Tissue sections from disease-free reduction mammoplasties (RMs) and five breast cancer subtypes (Luminal A (LumA), Luminal B (LumB), Her2-positive (Her2), triple negative (TNBC) and metaplastic (MBC)) were sequentially immunostained as described in Methods for *A*, neutrophil-enriched *myeloperoxidase* (MPO) detected with FITC, the myeloid marker *ITGAM/CD11b* detected with Cy3 and the immune cell marker CD45 detected with Cy5; or *B*, RNAse3/eosinophil-specific *Eosinophil cationic Protein* detected with FITC and the myeloid marker *ITGAM/CD11b* detected with Cy3. Stained slides were counterstained with DAPI and mounted prior to imaging. Representative images (at 20 x magnification) for one specimen of each of the 6 tissue specimen groups described above are shown. White arrows point to some of the numerous MPO-expressing neutrophils (panel *A*) or RNAse3-expressing eosinophils (panel *B*) in MBC. Scale bar: 0.1mm. C) Quantitative analysis of neutrophil myeloperoxidase (MPO) and eosinophil cationic protein (RNAse3). Images of immunostained breast tissue sections were subjected to quantitative analysis (Supplemental Table S5) as described in Experimental Procedures. Left panel: Numbers of MPO+ITGAM+ cells (neutrophils) per mm2 of analyzed tissue specimen are displayed for disease-free breast (RM; n=13) and each breast cancer subtype (LumA; n=20, LumB; n=9, Her2; n=8, TNBC; n=11 and MBC; n=8). Degrees of significance for each pair comparison (*, **) are displayed at the top of the panel. P-values: LumA vs. RM: p=0.0434; LumB vs RM: p=0.0203; Her2 vs. RM: p=0.0033; TNBC vs. RM: p=0.068; MBC vs. RM: p=0.0025. Right panel: Same as above for RNASE3+ITGAM+ cells (eosinophils). Same number of specimens as above except for TNBC; n=9. Degrees of significance for each pair comparison (*, **, ***) are displayed at the top of the panel. P-values: LumA vs. RM: p=0.0227; LumB vs RM: p=0.047; Her2 vs. RM: p=0.0022; TNBC vs. RM: p=0.023; MBC vs. RM: p=0.0008. Parameters used to generate graphs are shown in Supplemental Table S5.

The ’royal blue’ WGCNA module (Fig. 6*D*) encompassed proteins associated with unique aspects of MBC such as epigenetic plasticity, ectopic tissue formation, metastasis, tumor stiffness, and cellular survival under stress. The cartilage-associated protein (CRTAP) was upregulated predominantly in MBC (Supplemental Table S2), a subtype known for frequent ectopic bone tissue formation in the breast tumor and its propensity to metastasize to the bone (59). Interestingly, CRTAP has been implicated in the activation of Procollagen lysyl hydroxylase (PLOD) activities (60). Procollagen lysyl hydroxylase 2 (PLOD2) is a crucial player in the formation of fibrillar collagen in breast tissue and has been reported to regulate TGF-beta activity as well as enhance tumor stiffness (61), thereby promoting mesenchymal cancer cell phenotypes and stemness (62). PLOD2 upregulation has also been shown to correlate with tumors with poor outcome characterized by an extensive immune cell infiltration, in particular eosinophils, a distinct MBC phenotype as discussed below (63). Additionally, PLOD2, through interactions with alpha-ketoglutarate, can generate succinate which can subsequently suppress TET proteins and disrupt DNA demethylation. Another protein of interest was insulin-like growth factor binding protein-3 (IGFBP3), a protein known to stimulate survival and autophagy in breast cancer cells experiencing glucose deprivation and hypoxia (64), two common conditions in the tumor microenvironment. The collagen triple helix repeat-containing 1 (CTHRC1), a pro-metastatic protein, was found to have a positive correlation with the abundance of MMP7 and MMP9, thereby promoting tumor migration, motility, and invasion, particularly in non-small cell lung cancer (65). Furthermore, complement C1r subcomponent-like protein (C1RL), as part of the immune response, is activated during chronic inflammation (66).

The ‘green’ WGCNA module (Fig. 6*D*) was comprised of a diverse array of proteins with the highest upregulation in MBC. These proteins included the SH3 domain binding glutamate-rich protein like 3 (SH3BGRL3), which is involved in cell growth and apoptosis signaling pathways (67), and the quaking protein, KH domain RNA binding (QKI), which is crucial for post-transcriptional mRNA processing and cell differentiation (68). Pentraxin 3 (PTX3) is part of the innate immune system and is known to influence several aspects of cancer biology, including tumorigenesis and breast cancer metastasis to bone (69). The procollagen-lysine 2-oxoglutarate 5-dioxygenase 3 (PLOD3) is involved in collagen formation, a key component of the ECM, and is associated with tumorigenesis and metastasis (70). Cathepsin B (CTSB) is a cysteine protease involved in protein breakdown associated with various pathological states such as cancer and inflammation (71), and carboxypeptidase, vitellogenic-like (CPVL), contributes to protein processing and degradation. Of note, CPVL has been shown to facilitate resistance to CDK4/6 inhibitors in breast cancer (72). Furthermore, the simultaneous upregulation of CPVL, fatty acid binding protein 3 (FABP3), integrin subunit alpha 5 (ITGA5), orosomucoid 1 (ORM1), ORM2, ribosomal protein S19 (RPS19), and SERPINE2 and downregulation of transmembrane protein 205 (TMEM205) evoked heightened immunosuppression in MBC through polarization of macrophages toward a pro-tumorigenic M2 phenotype and/or through proliferation of regulatory T cells. Finally, the actin beta like 2 (ACTBL2) protein maintains cell structural integrity and facilitates processes such as motility and division (73).

### 3.11 | Metaplastic Breast Cancers (MBC) Display a Unique Tissue Landscape Typifying their Lineage Promiscuity

Strikingly, we discovered that MBC featured twice as many unique differentially changing signature proteins, in contrast to the other cancer subtypes that also showed unique signatures but to a much lesser extent, this was true even for TNBC. Overall, MBC featured 181 unique proteins that were upregulated and 168 unique proteins that were downregulated in MBC specifically (Fig. 2*F*). The identification of a large fraction of unique significant changes in protein levels in MBC (Supplemental Table S3-*B*) was anticipated (and now confirmed) based on the histological heterogeneity and the emergence of ectopic tissues or metaplasia (bone, muscle, etc.) in these rare, poorly differentiated, and particularly aggressive breast cancers (Supplemental Fig. S1). This unique plasticity in the breast for MBC was illustrated by the upregulation of previously defined ‘stemness’ proteins, such as adenosine monophosphate deaminase 2 (AMPD2), cluster of differentiation 109 (CD109), phosphoribosyl pyrophosphate amidotransferase (PPAT), and phosphoserine aminotransferase 1 (PSAT1), as well as downregulation of proteins, such as ATPase H+ transporting V0 subunit a1 (ATP6V0A1), a phenotype previously observed upon abnormal brain differentiation associated with lysosomal dysfunction(74) (Supplemental Fig. S3*C*). The suggested ‘stemness’ signature of MBCs was further exemplified by unique upregulation of proteins typically expressed in other tissues but not usually in breast tissue. Examples of such proteins included proteins reminiscent of osteocytes, i.e. bone-producing cells, such as osteolectin (CLEC11A). The upregulation of bone formation-associated proteins, specifically in MBC, aligned with its known propensity for ectopic bone (and cartilage) formation in the breast (75, 76) and bone dissemination (59). Additionally, MBC showed high keratin 17 (KRT17) level and anterior gradient 3 (AGR3) level, two phenotypes previously reported to be more frequent in TNBC and to be associated with worse outcome (77). This latter feature was likely an illustration of a squamous subtype of MBC. The striking upregulation of branched-chain-amino-acid aminotransferase (BCAT1) only observed in MBC (Supplemental Table S3*B*) further illustrated the complex cross-talk between epithelial cells and fibroblasts contributing to the acquisition of a CAF phenotype (78) and of a microenvironment favoring EMT and stemness (79). Finally, the extensive downregulation of selenium-binding protein SELENBP1 in MBC may also reflect a loss of cell differentiation. Previous reports demonstrate induction of SELENBP1 in colon cancer cells triggers colonocyte differentiation (80). A heightened EMT profile, characteristic of tissues with high plasticity, and previously reported as a major phenotype in MBC (5), could be validated in our FFPE specimen analysis. Indeed, 290 of the 1153 ‘EMTome’ proteins (81) were captured in our proteomic analysis. Seventeen proteins were upregulated and 13 proteins downregulated specifically in MBC. The most consistent of these MBC-specific changes included upregulation of CD44, NDRG1, SERPINE2 and TGFBI and downregulation of AP1M2, SELENBP1 and STAT5A. The detection of such a wide array of unexpected proteins in breast, unique to MBC, highlighted the exquisite adaptability of cancer cells in a quite unique tumor microenvironment.

### 3.12 | MBC: Extensive Deposition of Proteins Reflecting Chronic Inflammation and Involvement of Coagulation

Besides upregulated proteins involved in cell proliferation, cell survival and cell migration/invasiveness biological processes, the MBC-specific signature identified many upregulated proteins involved in chronic inflammation, fibrosis, and immune response (Supplemental Fig. S7, *A* and *B*). Thus, we observed upregulation of a panel of proteins indicative of chronic inflammation (Supplemental Table S3*B*): serum amyloid A-4 protein (SAA4), alpha-2-HS-glycoprotein (AHSG), tumor necrosis factor alpha-induced protein 8 (TNFAIP8), transthyretin (TTR), typically expressed in liver, choroid plexus, pancreas and kidney, leukosialin (SPN), typically expressed in lung, skin and bone marrow, plasminogen (PLG) typically expressed in liver and kidney; complement proteins (C4BPA, C4BPB and C8B) typically expressed in liver, lung and/or, testis; and afamin (AFM) typically expressed in liver, testis and kidney. Upregulation of afamin (AFM) in MBC strongly supported stabilization of Wnt ligands and as a result heightened Wnt signaling through this unique mechanism. The upregulation of transforming growth factor-beta-induced protein ig-h3 (TGFBI) also reflected engagement of a key node in fibrosis pathways. Additional markers typifying (chronic) inflammation included upregulation of agrin (AGRN), interleukin 18 (IL18), protein arginine methyltransferase 1 (PRMT1), SERPINA3, SERPINA4, SERPINB8, SERPINB9, SERPIND1, SERPINF2, and Thimet oligopeptidase 1 (THOP1), as well as downregulation of repressors of inflammation and fibrosis, such as mediator complex subunit 23 (MED23). It was intriguing to observe that MBC-specific protein signatures were indicative of activation of complement (C) or coagulation factors (F), as illustrated by upregulation of C7, CFI, F2, F9, F12, F13A1, kininogen 1 (KNG1), plasminogen (PLG), a biological function previously reported as enriched in MBC (5). It has recently emerged that activation of this coagulation cascade is not only leading to platelet, neutrophil and endothelial cell activation but also promotes cancer cell growth and invasion (82).

A significant number of the protein level changes observed may be attributable to increased MYC activity. Indeed, a myc signature has been identified in MBC (5) but also in (metaplastic) lung squamous cell carcinoma but not in (non-metaplastic) lung adenocarcinoma (83). A systematic search of the proteins captured in our proteomic analysis among a 198 ‘HALLMARK MYC TARGET’ published database (84) identified 122 ‘hallmark myc target’ proteins. Five of these proteins were upregulated (EIF3I, HNRNPA2B1, PPIA, PSMA1 and UBA2) and 1 protein downregulated (CSTF2) specifically in MBC. The most consistent MBC-specific change was the upregulation of peptidylprolyl Isomerase A (PPIA).

### 3.13 | Breast Tissue Injury and Aging leads to Dysregulation of Stromal Proteins and Cancer Progression

Cancer progression and tumorigenesis is facilitated and perhaps even driven by disruptions to the ECM that lead to the unrestrained release of soluble factors and changes in mechanotransduction, thus creating a chronically inflamed microenvironment that favors cancer progression and increased senescence burden (Fig. 4*A*). A graphical display of MBC vs disease-free significant protein changes was visualized using a map of proteins localized to organelles (Supplemental Fig. S7*C*) (85, 86). This display indicated significant changes in levels of proteins involved in regulation of mitochondria, peroxisome, endoplasmic reticulum, and proteasome function. A predominant downregulation of mitochondrial and peroxisomal proteins would support reduced energy production (87, 88), while upregulation of endoplasmic reticulum and proteasome proteins would correlate with increased signaling and demand to degrade proteins as the cells enter a stress mode associated with tumorigenesis (89, 90). These changes, specifically observed for MBC, highlighted characteristics of a disrupted tissue microenvironment in metaplastic cancers that might contribute to the unique lineage promiscuity and cell plasticity, and importantly, to the heightened aggressiveness of these rare cancers. In our investigation of the coping mechanisms of cancer cells amidst various neoplastic stresses, we observed notable alterations in endoplasmic reticulum (ER) proteostasis (Fig. 4*B* and Supplemental Fig. S5*B*). Proteostasis relies on regulated protein translation, proper protein folding by molecular chaperones, and activation of protein degradation pathways such as autophagy and proteasome-mediated degradation. The fine tuning of these mechanisms in response to homeostatic variations or stress conditions is pivotal to the maintenance of an appropriately folded proteome. This tuning relies on proper and timely engagement of cellular stress response pathways -heat shock response (HSR) as well as mitochondrial and ER unfolded protein responses (UPRs) (91). This is a dual-edged phenomenon that can promote cell adaptation and survival under moderate stress but initiate instead cell suicide under persistent stress (89). The upregulation of various ER proteins in breast cancer subtypes (Fig. 4*B*) compared to disease-free tissue underscores the importance of ER proteostasis in neoplastic stress responses, highlighting the importance of ER-based processes in both the promotion and suppression of tumorigenesis. The upregulation of flap endonuclease 1 (FEN1), in conjunction with the downregulation of double strand break repair protein (RAD50, the most downregulated protein in the MBC dataset; -6.15) and of inosine triphosphatase (ITPA), further illustrated profound defects in DNA damage repair expected in stressed tissues upon injury or aging.(92)

## 4 | DISCUSSION

The generation of these in-depth comparative proteomic profiles of five major human breast cancer subtypes and disease-free human breasts was the result of the use of modern mass spectrometry technology in combination with the distinctly optimized FFPE sample processing through extensive sample deparaffinization and de-lipidation (Folch Extraction), that became necessary due to the high adiposity of breast tissues. The optimization of an efficient quantitative proteomic workflows for FFPE cancer tissues opens access to extensive archived FFPE tissues from biobanks. This becomes particularly relevant when conducting longitudinal studies using retrospective cohorts with long term patient follow-up to assess, for example, disease progression or relapse for risk stratification purposes or response to therapy. In this study, we were able to decipher acellular tissue components (ECM) without ECM enrichment strategy and together with the cellular tissue components we were able to query a wide range of biological phenotypes. Furthermore, while some of our observations confirmed and extended epithelial phenotypes during breast cancer, they also extended our insights into key stromal phenotypes, both in terms of cellular and acellular tissue components, i.e. stromal cells and ECM, respectively.

Thus, we have delineated across all breast cancer subtypes conserved upregulation and downregulation of proteins that support: (1) increased proliferation typified by higher levels of proteins involved in transcription and translation, (2) switches in lipid and sugar metabolism as well as mitochondrial function, and (3) extensive tissue remodeling associated with stress response and differentiation. Although some of these changes were shared among all subtypes, a significant number of them showed a gradual increase from more differentiated breast cancers (luminal A and B) to less differentiated breast cancers (TNBC and MBC). This observation afforded us with the ability to highlight malignant events whose extent of dysregulation seemed to correlate with tumor aggressiveness. A more granular analysis of the various breast cancer subtypes identified protein signatures unique to each subtype. The least differentiated breast cancer, MBC, displayed several striking phenotypes.

First, in MBC, we observed a remodeling of ECM illustrated by an extensive loss of ECM proteins: downregulation of collagens, in particular collagens 4, 6, 15 and 18, as well as small leucine rich protein decorin. A loss of desmosome proteins (desmocollins, desmogleins and desmoplakin) illustrated loss of cell-cell contacts that was going hand in hand with an extensive EMT signature. Indeed, just as striking was the upregulation of ECM proteins known to foster a pro-tumorigenic environment including periostin, fibronectin, versican and tenascin (93–95). The upregulation of fibromodulin, specifically restricted to MBC, was noteworthy as its upregulation has been linked to breast cancer metastasis (96). Furthermore, this phenotype is in accordance with increased Wnt-beta catenin signaling and is a mark of chronically inflamed tissue. The upregulation of fibromodulin can be inhibited by anti-inflammatory drugs such as aspirin. Notably, we observed a specific upregulation of pivotal ECM-modifying proteins, such as MMP9, a protease widely implicated in tumorigenesis, as well as neutrophil collagenase MMP8. The upregulation of such ECM-proteolytic enzymes specific to MBC had far reaching consequences, not only leading to an extensive loss of ECM architecture but also more broadly contributing to a loss of tissue homeostasis and to the acquisition of a particularly aggressive and immunosuppressed tumor microenvironment. The upregulation of PLOD proteins, maximal for PLOD1 and PLOD3 and unique for PLOD2 in MBC, and the ensuing extensive collagenolysis have been recently related to promoting desmoplastic cancers with very poor patient outcome (63). The upregulation of BCAT1 was another example of an epithelium-stroma cross-talk leading to acquisition of a CAF phenotype associated with cancers with heightened EMT and cancer cell stemness (78, 79).

Our analysis identified profound changes affecting multiple stromal tissue components. For example, the loss of ADH1B illustrated a shift in fibroblast identity towards a CAF phenotype (26). The loss of CD36, previously reported by us across all breast cancer subtypes (27), reflected a shift in vasculature identity characterized by a loss of physiological vasculature architecture and identified by the detachment of a recently characterized capillary population involving pericytes (29). The upregulation of myosin 9 (MYH9) and the myofibroblast-specific myosin 11 (MYH11) further exemplified the acquisition of a highly EMT phenotype in breast cancers (97). Lastly, the loss of several mast cell-specific proteins (chymase (CMA1) and carboxypeptidase A (CPA3)) further documented these changes in tissue ecology. These relationships unveil the too long ignored proclivity of an altered stroma to dictate epithelial tumor cell fate. We document below the remarkable changes observed in immune tissue landscape.

The importance of the immune (stromal) environment first indicated by Finak et al. (8), has revolutionized classification of tumors and associated patient outcome. Highlighting the importance of this tissue ecology aspect, our second observation was that MBC was characterized by an immunosuppressed tumor environment. Indeed, while alterations in immune response were seen across all 5 subtypes of breast cancer, this phenotype exhibited the most variation that was unique to each subtype. The concomitant upregulation of EPX, RNASE3, DEFA1, MPO, PRTN3, ELANE and ITGAM, identified through proteomic analysis (Supplemental Table S3-*B*) and WGCNA (Fig. 6*D*), documented recruitment of eosinophils and neutrophils, as a phenotype exemplified in the context of the MBC tissue microenvironment (98, 99). Due to its unique mode of action relying on the use of hydrogen peroxide to form different hypohalous acids leading to the production of reactive oxygen and reactive nitrogen species, uncontrolled or sustained release of MPO would exaggerate inflammation and contribute to oxidative stress-associated tissue damage even in absence of inflammation (100). We have previously reported upregulation of such biomarkers in other human CIACs, i.e., esophageal (10) and lung (14). It therefore appears that recruitment of specialized immune cells is a prominent phenotype of CIACs shared across tissue types, i.e., metaplastic breast, lung, and esophageal cancers. We have also recently identified eosinophil and neutrophil signatures (EPX and MPO upregulation) in a mouse model of colitis, highlighting the importance of such biological changes as early events in chronic inflammation-driven tumorigenesis (101). Previous findings in oral cancer models directly linked tumor-associated neutrophils to metastatic spread (57). Furthermore, neutrophils have been shown to play a key role in maintenance of metaplastic phenotypes in a murine model of lung metaplastic carcinoma, i.e. lung squamous cell carcinoma (9). Relevant to this study, MPO-derived oxidants have been involved in inducing premature cell senescence (102). Additional MBC-specific proteomic signatures further evoked heightened immunosuppression through polarization of macrophages toward a pro-tumorigenic M2 phenotype (typified by a loss of CD36 level) and/or through proliferation of regulatory T cells. Of particular novelty and introducing a new concept, recent evidence has pointed to the activation of neutrophil extracellular traps by SASP factors, which further promotes cancer cell proliferation, invasion, and metastasis in contrast to the traditional view of neutrophils killing pathological cells (103, 104). Our data further supported this hypothesis as SASP and neutrophilic proteins that give rise to neutrophil extracellular traps showed the highest abundance in MBC.

Third, another major trend was the more extensive deposition of proteins reminiscent of chronic inflammation, a key driver of tumorigenesis for metaplastic cancers. Indeed, we observed upregulation of a large panel of proteins indicative of chronic inflammation (AGRN, AHSG, IL18, PRMT1, SAA4, SPN, TNFAIP8, TTR) and complement proteins (C4BPA, C4BPB and C8B). The upregulation of some of these proteins suggested dysregulated signaling: sustained Wnt signaling associated with AFM upregulation or TGF-beta signaling associated with upregulation of TGFBI. A striking observation was the upregulation of serpins. We extensively discussed the upregulation of SERPINH1, but other serpins were upregulated as well. This pro-inflammatory signature was exemplified by the downregulation of repressors of inflammation and fibrosis (MED23). An intriguing observation was a MBC-specific protein signature indicative of deposition of proteins involved in activation of complement (C7, CFI) or coagulation factors (F2, F12), kininogen 1 (KNG1) and plasminogen (PLG). As discussed above, activation of this coagulation cascade is not only leading to platelet, neutrophil and endothelial cell activation but also contributes to cancer cell growth and invasion (82).

Lastly, our study extended the recent concept that (breast) cancer evolves as an accelerated process of tissue aging (105, 106). Such ‘biological’ aging likely results from a response to stress exemplified by chronic inflammation driving tumorigenesis in the case of metaplastic cancers. Aging-associated alterations in the ECM architecture and remodeling include loss of basement membrane components, increased collagen cross-linking and stiffness. We detected an upregulation of a large number of proteins associated with a senescence (SASP) phenotype (38). The risk and incidence of breast cancer increases with age (1), as does the accumulation of senescent cells, i.e., of cells that are in a state of permanent cell cycle arrest (107). Senescent cells are known to secrete inflammatory molecules and cytokines into the ECM (38, 108), where these senescent associated secretory phenotype (SASP) factors promote increased tissue stiffness and a tumor-prone microenvironment (108). In our study, activation of senescence and inflammatory pathways was evident across breast cancer subtypes through conserved upregulation of numerous SASP factors (Fig. 4*C*). Further extending a stress response signature in breast cancers, we observed notable alterations in endoplasmic reticulum (ER) proteostasis and an upregulation of many damage-associated molecular patterns (DAMPs) including ECM proteins (VCAN, TNC) but also several heat shock proteins (HSPA1A, HSPA5, HSPA8, HSP90AA1, HSP90AB1 and HSP90B1). Upregulation of one of these DAMPs, CD44, along with SASP-Associated candidate, SERPINF2 was only observed in MBC. The consistent overexpression of proteins like SERPINH1, POSTN, TNC, and FN1 across breast cancer subtypes, also implicated as pro-tumorigenic drivers in LSCC (14), implied the common engagement of local and systemic SASP programs as one of the programs promoting cancer progression. Relevant to this discussion, engagement of such proteins influences not only immune function but also ECM processing (109).

The resulting cell secretome likely acts both locally to remodel the ECM and distally to spread inflammatory signals, thus contributing to sustained matrix stiffening, chronic inflammation and more broadly a tumor-supportive microenvironment. The role of ECM alterations is also closely related with aging, as ECM stiffening with age presents additional prevalence for cancer invasion (32). Changes in tissue (stromal) architecture can already be identified at a pre-malignant stage and the extent of these changes can be used as a tool for risk stratification for developing subsequent breast cancer (29). Interestingly, in many breast cancers, epithelial cells show a striking loss of lineage fidelity. Thus, luminal epithelial cells start to lose their lineage fidelity, and begin to express markers typically associated with myoepithelial cells (106). This fidelity to cellular identity and function is compromised during the aging process and cancer development, as DNA damage incites inflammation and disruption of tissue homeostasis (105). Inflammatory processes gradually undermine the homeostatic cues maintaining cellular identity, leading to loss of cellular differentiation and further disarray in cellular function.

## 4 | CONCLUDING REMARKS

Our findings provide a foundation to further elucidate mechanistic events linking loss of cell identity and function observed in cancer to chronic inflammation, stress response and accelerated (chronological) aging (105). Knowledge of the molecular and pathological differences between the various breast cancer subtypes studied here, not only focusing on the epithelial component but also taking into consideration cellular and acellular stromal components, is critical for the development of therapies customized to each breast cancer subtype. However, we also note as limitation of our study the relatively small cohort size (although it should be noted, that for the very rare MCP it is not easy to obtain tissues). We consider this a first Pilot Study and additional patient cohorts will be needed to further validate our findings. Interestingly, despite the relatively small human cohort size, distinct protein signatures emerged throughout all cancer subtypes – common and unique – with remarkable statistical significance for many cancer-relevant proteins and pathways. Moving forward, high-priority foci for future work include: i) immunohistochemistry-based validation of differential levels of subtype protein markers in larger/independent cohorts, ii) elucidation of mechanisms via functional studies, and iii) exploration of biomarker applications to design therapeutic regimens targeting some of the pathways highlighted in this study. The development of subtype-specific protein signatures into clinically viable tests could enable better prognostication and therapy selection. With further mechanistic clarification and clinical validation, our proteomic findings may ultimately translate into personalized management strategies tailored to each patient’s tumor profile.

## ACKNOWLEDGMENTS

FFPE tissue blocks used for proteomic and mIHC analyses were obtained from the Cooperative Human Tissue Network Western and Mid-Atlantic divisions. The authors acknowledge the very generous support from SCIEX for the ZenoTOF 7600 system and a Waters M-class HPLC system at the Buck Institute.

## DATA AVAILABILITY STATEMENT

The raw data and complete mass spectrometry data sets have been uploaded to the Mass Spectrometry Interactive Virtual Environment (MassIVE) repository, which is maintained by the Center for Computational Mass Spectrometry at the University of California San Diego. These data can be accessed and downloaded using the following link: https://massive.ucsd.edu/ProteoSAFe/dataset.jsp?task=bfbb9bbd82244bd0810b6c5809 <u>db91ca</u> (MassIVE ID number: MSV000094056; ProteomeXchange ID: PXD049314). The R scripts used in this study are available through our GitHub repository (https://github.com/JoBBurt/Human-FFPE-Breast-Cancer-Subtypes) and can be cited using the DOI: https://doi.org/10.5281/zenodo.10651533. [Note to the reviewers: To access the data repository MassIVE (UCSD) for MS data, please use: Username: MSV000094056_reviewer; Password: winter].

### Supplemental data – This article contains supplemental data

*Author contributions* – J.B.B., B.S., P.G. and T.D.T. manuscript writing; J.B.B., D.P., R.B., C.C.-T. and J.A.C. methodology; J.B.B. and J.B. investigation; J.B.B., J.B. and B.S., formal analysis; J.B.B., B.S. and P.G. data curation; P.G., B.S., and T.D.T conceptualization; B.S. and T.D.T. supervision; C.L.H, B.S. and T.D.T. resources; P.G. project administration; B.S., P.G. and T.D.T. funding acquisition.

## Abbreviations

DIA: data-independent acquisition
ECM: extracellular matrix
ER: Estrogen Receptor
FDR: false discovery rate
FFPE: Formalin-Fixed Paraffin-Embedded
GO: gene ontology
Her2: human epidermal growth factor receptor 2
LumA: Luminal A
LumB: Luminal B
MBC: metaplastic breast cancers
MS/MS: tandem MS
mIHC: Multiplex Immunohistochemistry
PR: Progesterone Receptor
RM: reduction mammoplasty
SASP: Senescence-Associated Secretory Phenotype
TNBC: triple negative breast cancer

