## Supplementary material for "Proteomic Analysis of Breast Cancer Subtypes Identifies Stromal Protein Profiles that Contribute to Aggressive Malignant Behavior": Suppl Figure 1

### Supplementary Figure S1

A **Lesions with Mixed Histology in Metaplastic Breast Cancer Tissue Specimens**

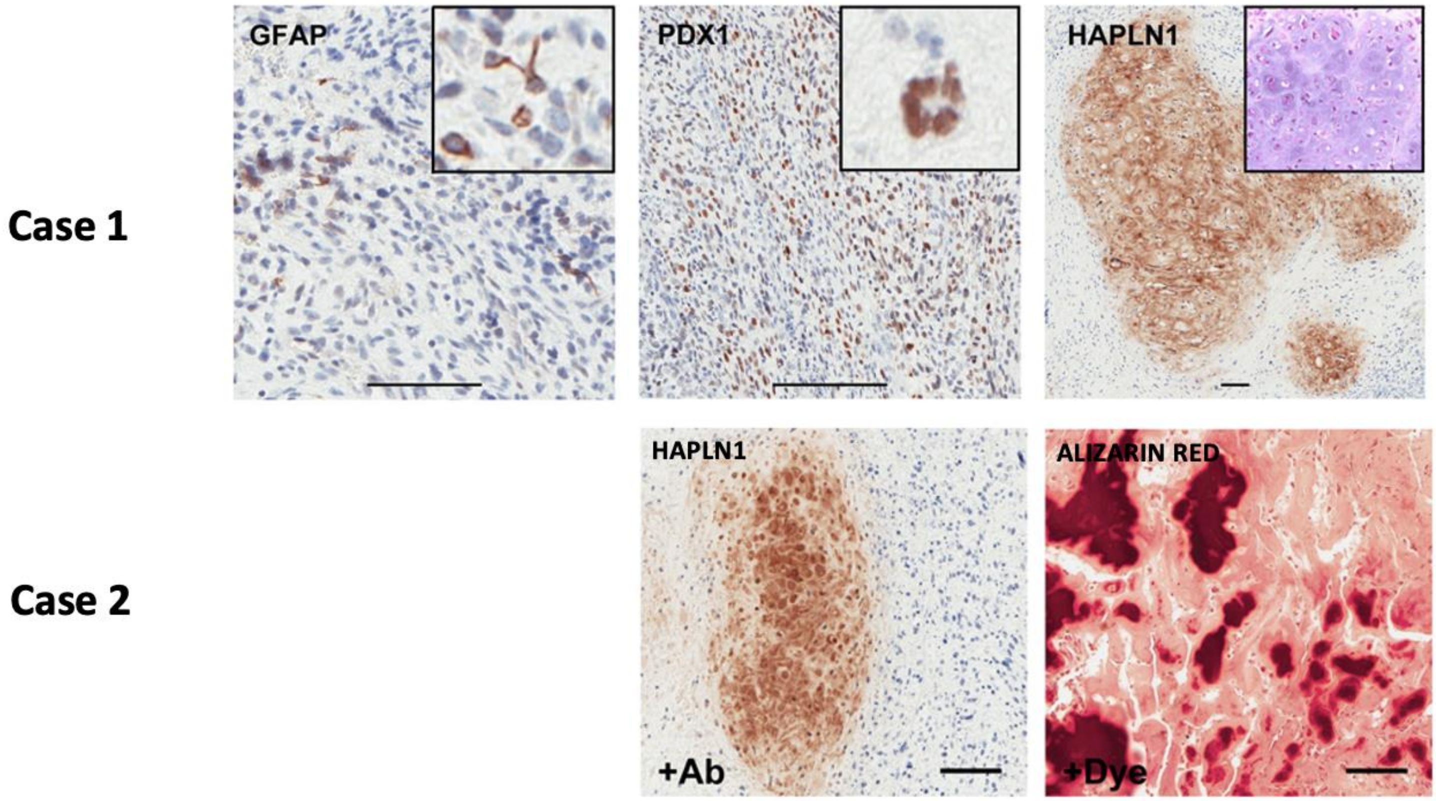

### Supplementary Figure S1

B **Histology of Disease-free Breast Tissue Specimens**

$B_1$ **RM 175**

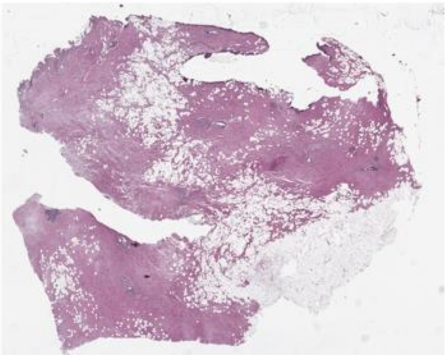

$B_2$ **RM 94**

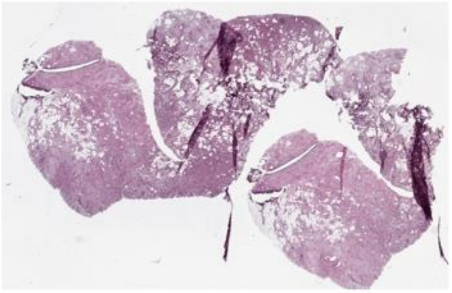

$B_3$ **RM 106**

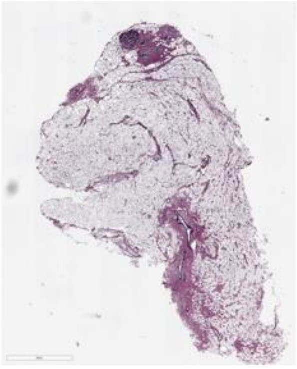

$B_4$ **RM 291**

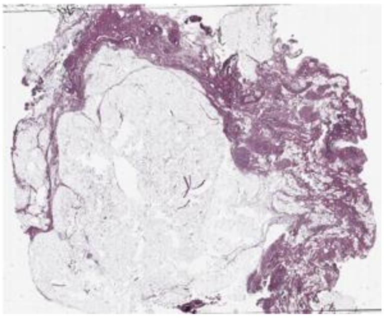

$B_5$ **RM 95**

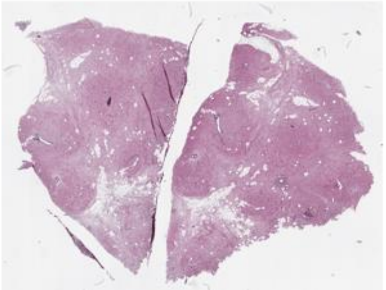

$B_6$ **RM 104**

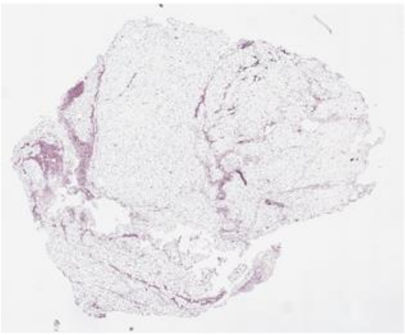

$B_7$ **RM 117**

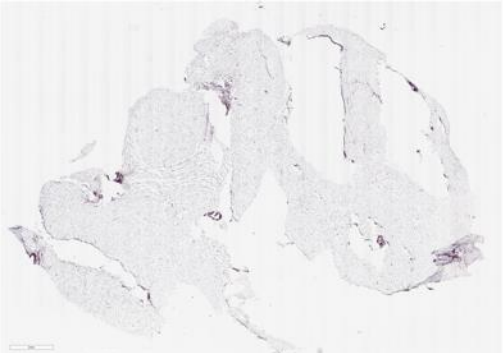

### Supplementary Figure S1

#### C Histology of Luminal A Breast Cancer Tissue Specimens

$C_1$ **WD-80873**

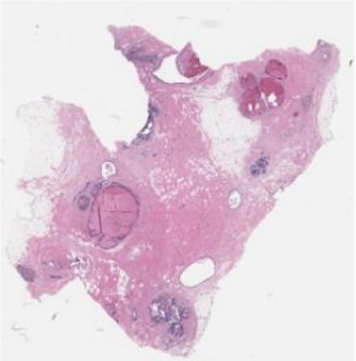

$C_2$ **WD-80874**

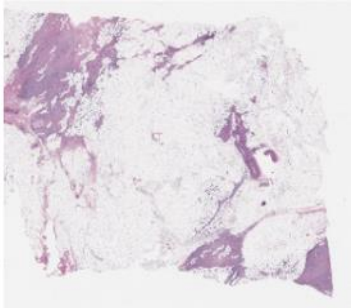

$C_3$ **WD-80876**

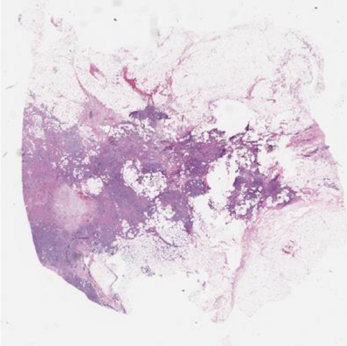

$C_4$ **WD-80877**

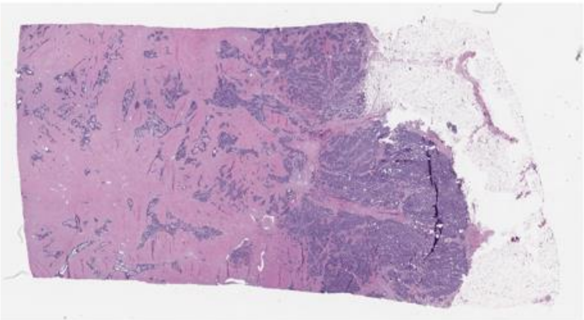

$C_5$ **WD-80878**

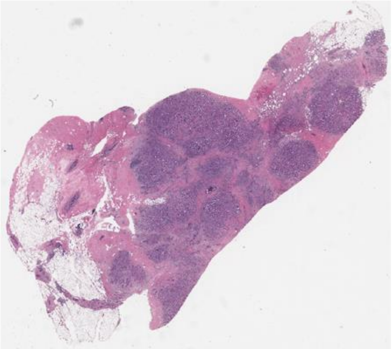

$C_6$ **WD-80879**

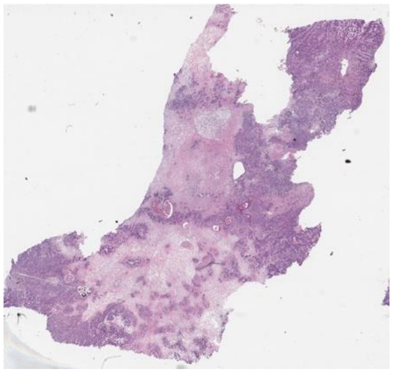

$C_7$ **WD-80880**

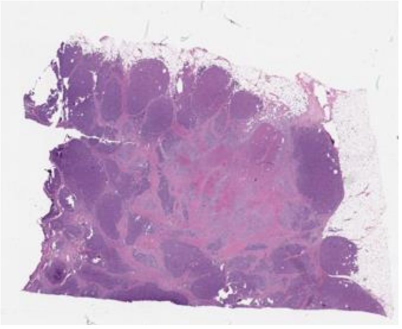

### Supplementary Figure S1

D **Histology of Luminal B Breast Cancer Tissue Specimens**

$D_1$ **WD-80884**

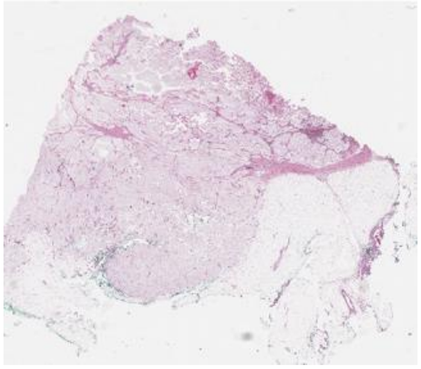

$D_2$ **WD-80886**

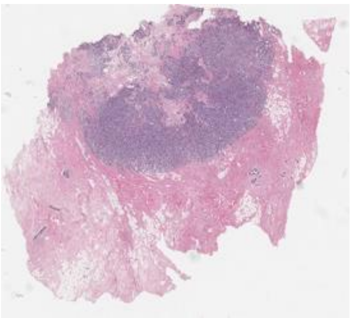

$D_3$ **WD-80887**

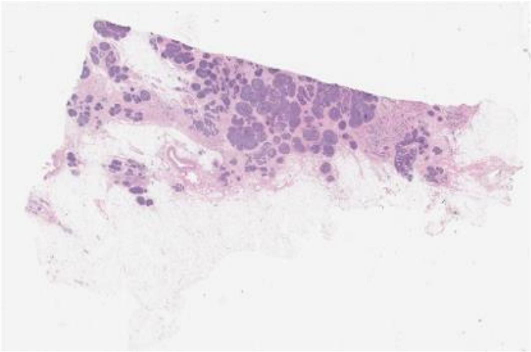

$D_4$ **WD-80889**

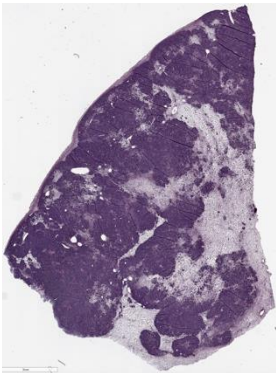

$D_5$ **WD-80894**

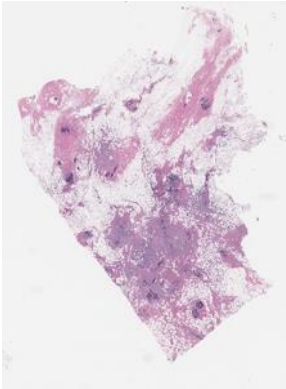

$D_6$ **WD-80896**

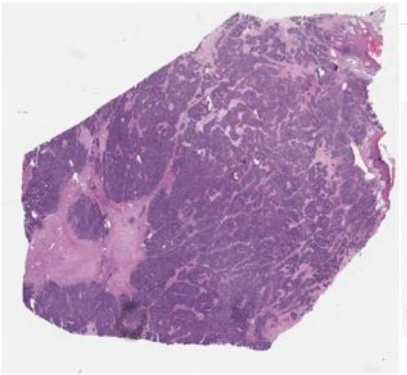

$D_7$ **WD-80897**

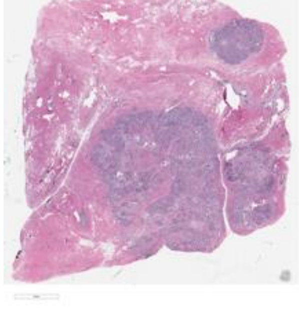

### Supplementary Figure S1

E **Histology of Her2+ Breast Cancer Tissue Specimens**

$E_1$ **WD-80872**

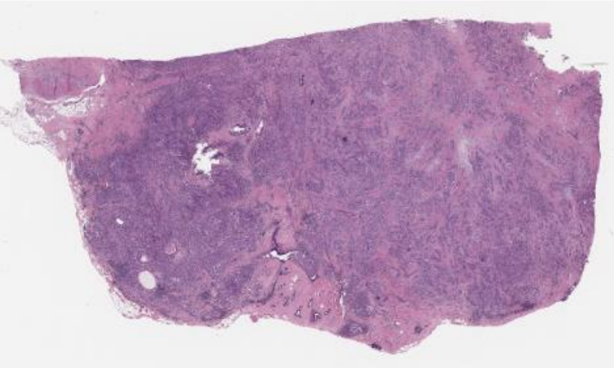

$E_2$ **WD-80888**

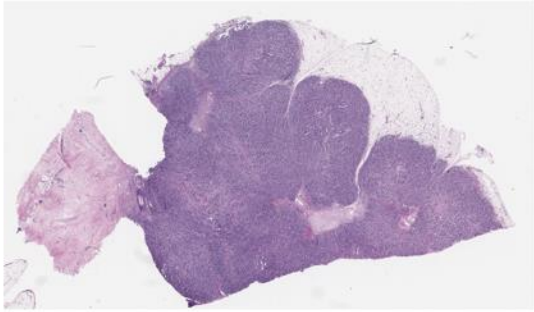

$E_3$ **WD-80890**

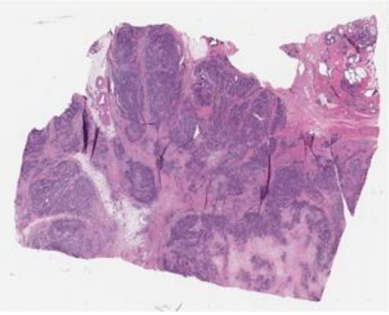

$E_4$ **WD-80891**

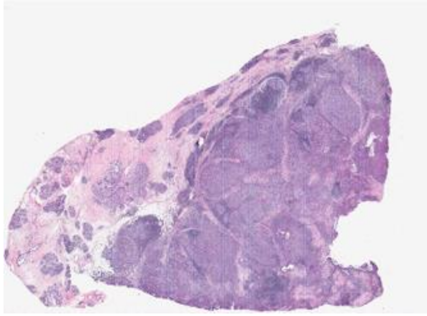

$E_5$ **WD-80892**

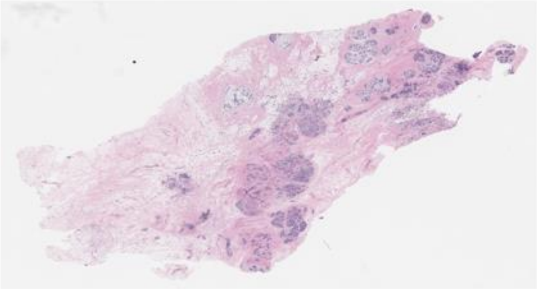

$E_6$ **WD-80900**

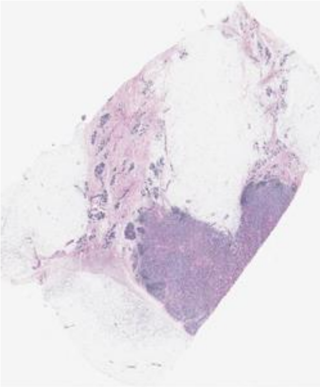

$E_7$ **WD-80901**

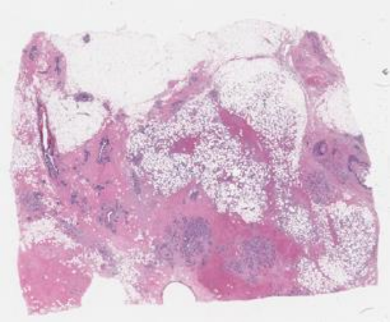

### Supplementary Figure S1

F **Histology of Triple Negative Breast Cancer Tissue Specimens**

$F_1$ **WD-80875**

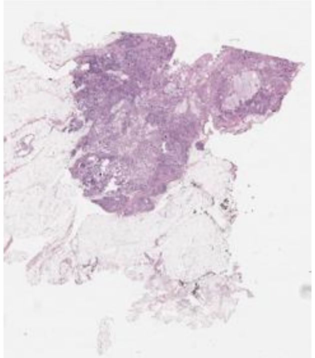

$F_2$ **WD-80885**

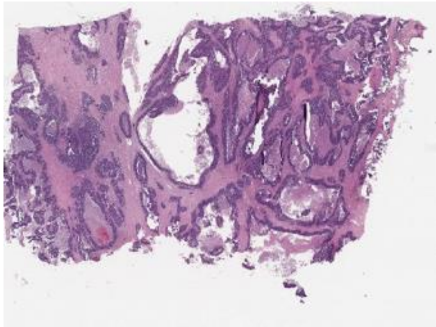

$F_3$ **WD-80893**

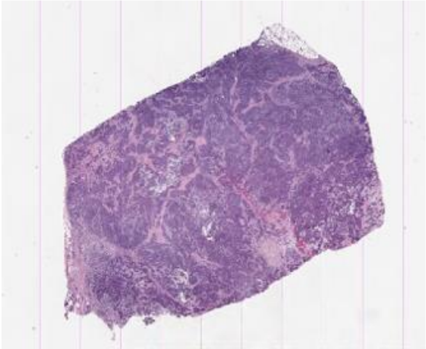

$F_4$ **WD-80895**

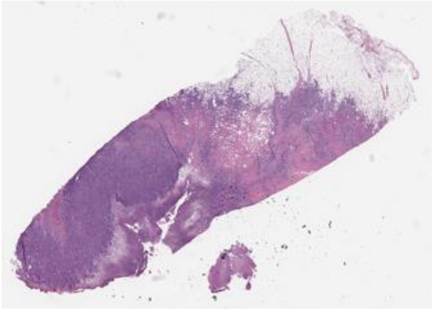

$F_5$ **WD-80902**

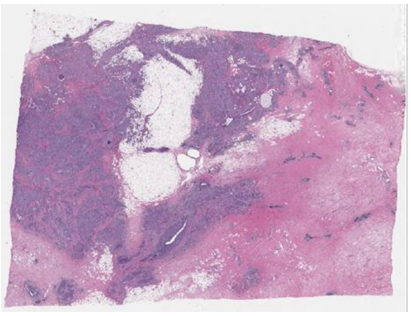

$F_6$ **WD-80903**

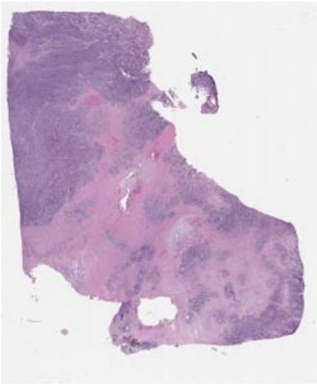

$F_7$ **WD-80904**

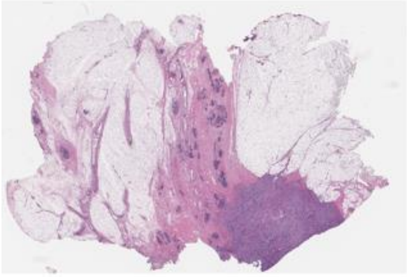

### Supplementary Figure S1

#### G Histology of Metaplastic Breast Cancer Tissue Specimens

$G_1$ **WD-82424**

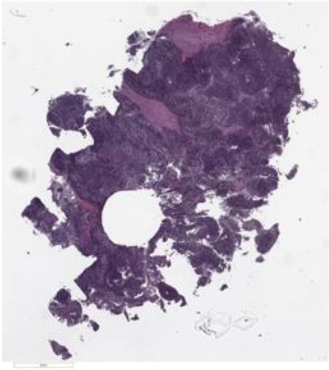

$G_2$ **69928T001**

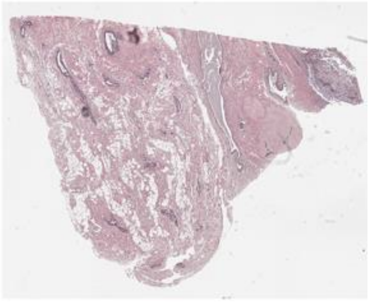

$G_3$ **89165T001**

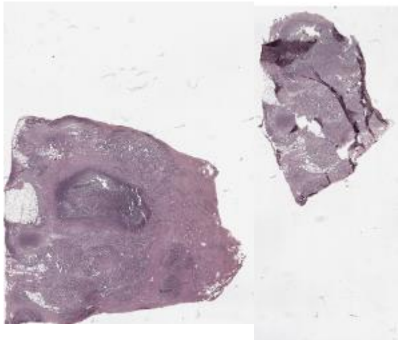

$G_4$ **89789T002**

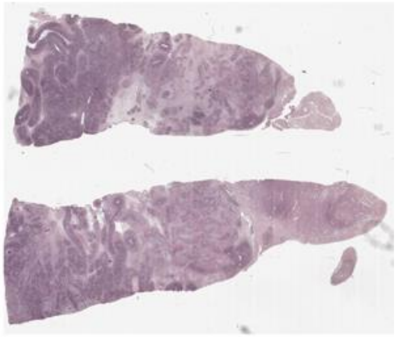

$G_5$ **MAD09-00008**

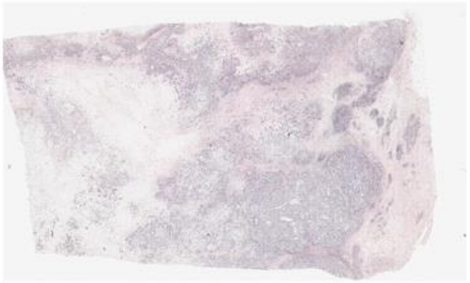

$G_6$ **MAD11-00398**

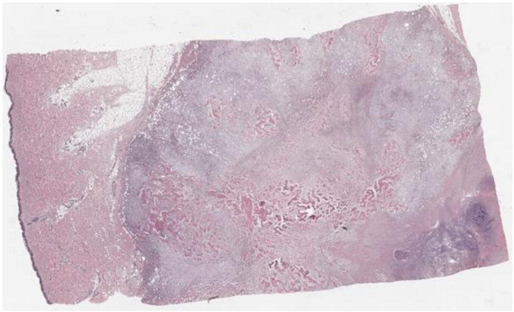

$G_7$ **MAD11-00715**

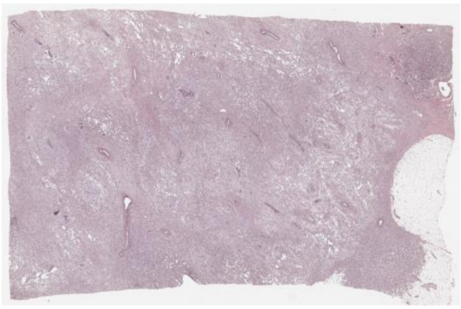

**Supplementary Figure S1. Histology of Disease-free Breast and Breast Cancer FFPE Tissue Specimens.** Immunohistochemical stainings of tissue sections from two cases of metaplastic breast cancer are shown in panel A. Upper row (case 1): evidence of breast lesions with ectopic expression of the neuronal-specific marker GFAP, pancreatic-specific marker PDX1 and cartilage-specific marker HAPLN1 revealed with DAB (brown dye). Lower row (case 2): ): evidence of breast lesions with ectopic expression of the cartilage-specific marker HAPLN1 revealed with DAB (brown dye) and bone revealed with Alizarin Red (red dye) . Hematoxylin and eosin stained FFPE tissue sections are shown for B) disease-free reduction mammoplasty (RM), C) Luminal A (LumA), D) Luminal B (LumB), E) HER2-positive (HER2), F) triple negative (TNBC), and G) metaplastic (MBC) breast tissue specimens used in this study.
