## Supplementary material for "Proteomic Analysis of Breast Cancer Subtypes Identifies Stromal Protein Profiles that Contribute to Aggressive Malignant Behavior": Suppl Figure 2

### Supplementary Figure S2

##### Gene Ontology of All Identified Proteins

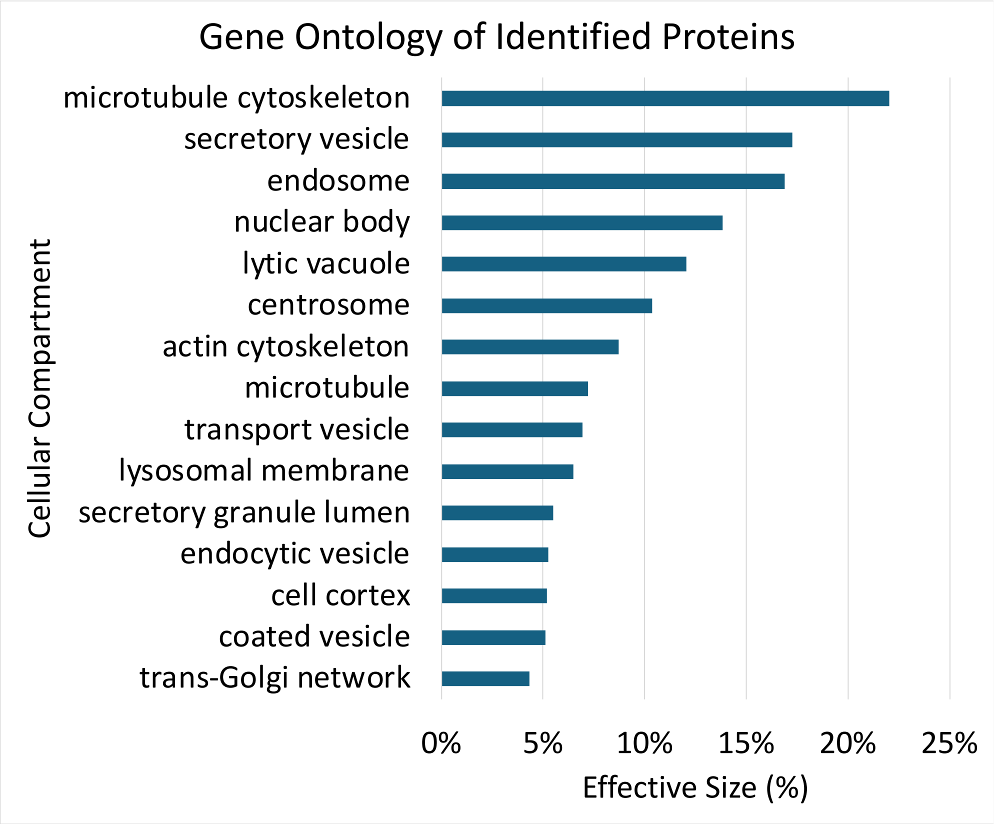


**Supplementary Figure S2. Cellular compartment representation among proteins with altered expression in breast cancer subtypes compared with disease-free tissue.**

Barplot of KEGG cellular compartments from all proteins identified in the dataset, the effective size is the number of proteins identified in each compartment divided by the total number of proteins identified. The displayed KEGG biological processes, molecular functions, and cellular compartments have a q-value ≤ 0.01 and term level ≥ 5.
