## Supplementary material for "Proteomic Analysis of Breast Cancer Subtypes Identifies Stromal Protein Profiles that Contribute to Aggressive Malignant Behavior": Suppl Figure 3

### Supplementary Figure S3

A **Ratios for 576 Significantly Altered Protein Groups in all Subtypes vs Control**

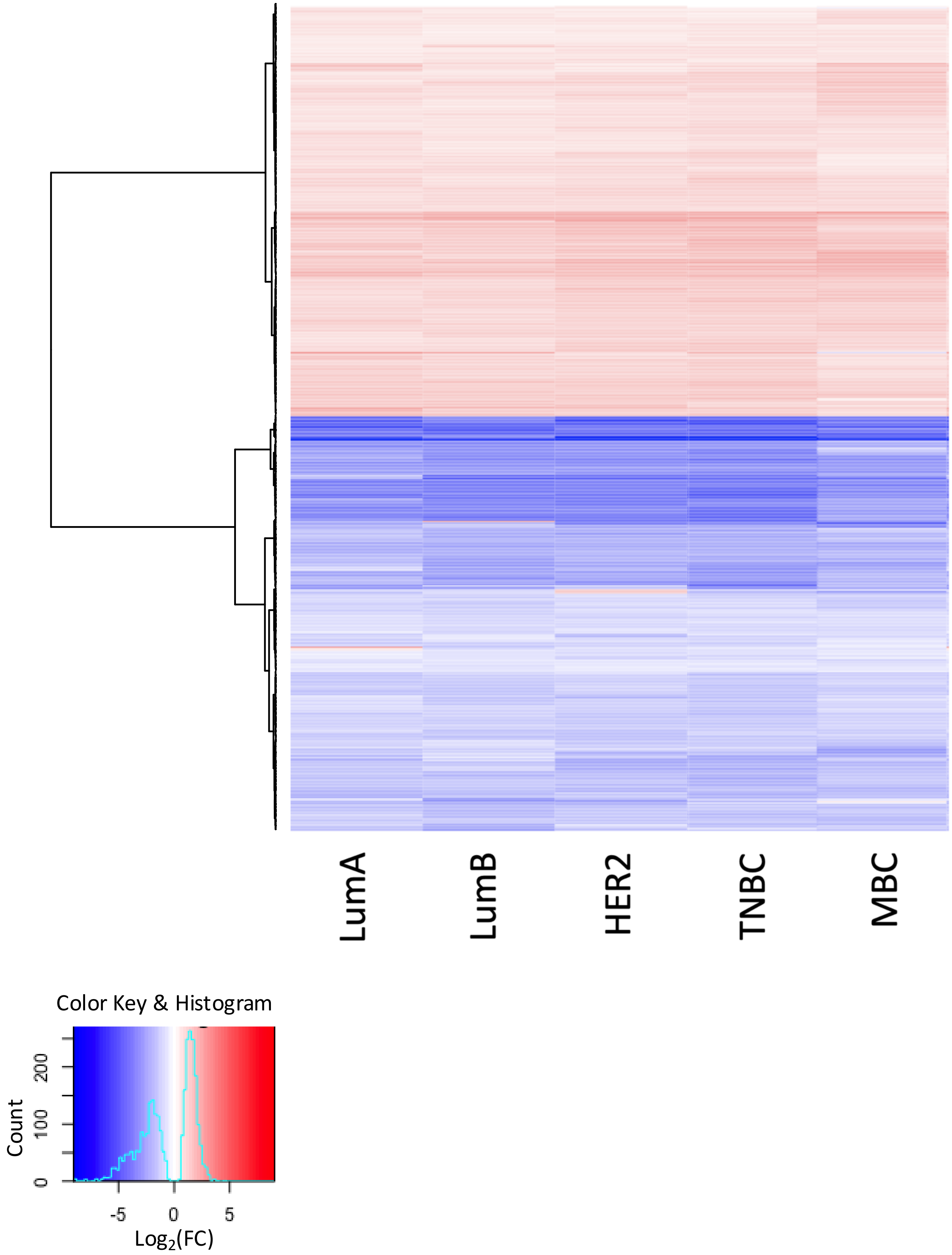

### Supplemental Figure S3

B **Protein Groups with Fold Changes that Reflect Breast Cancer Subtype**

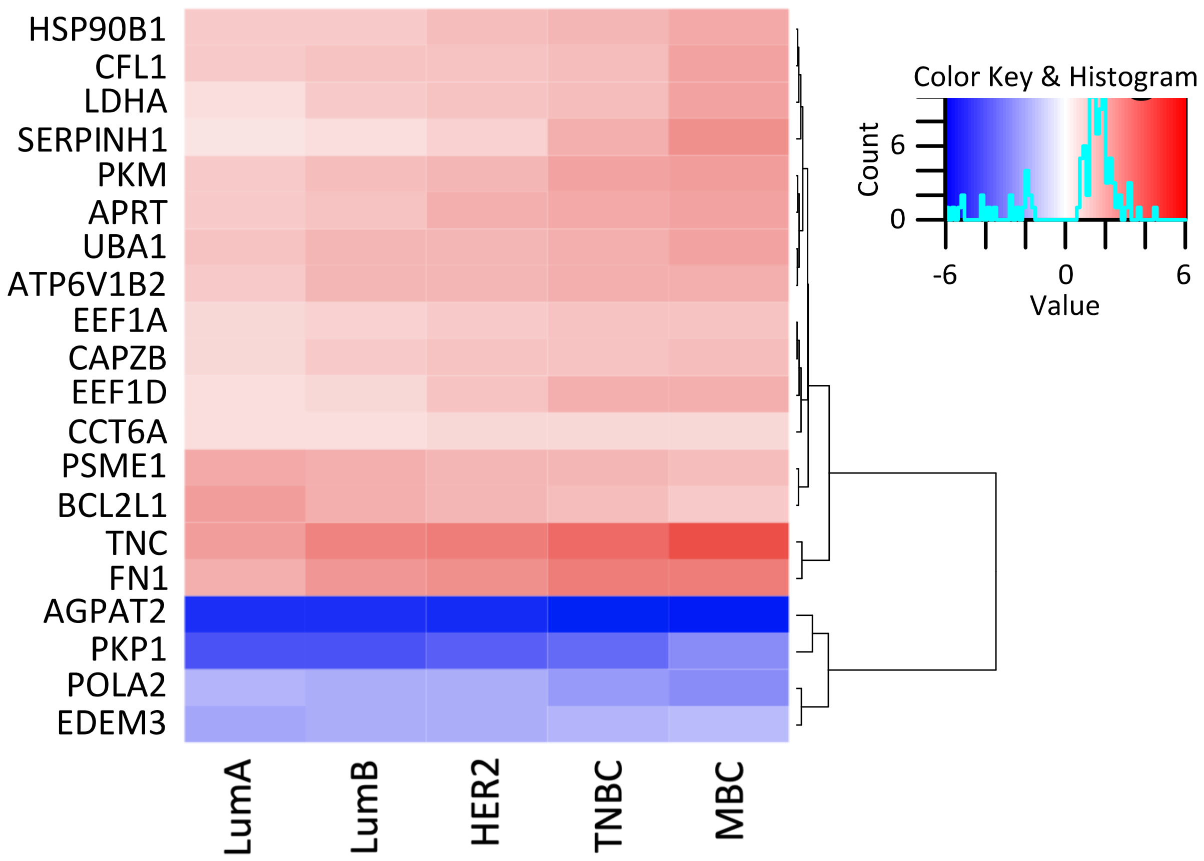


C **Stemness Markers**

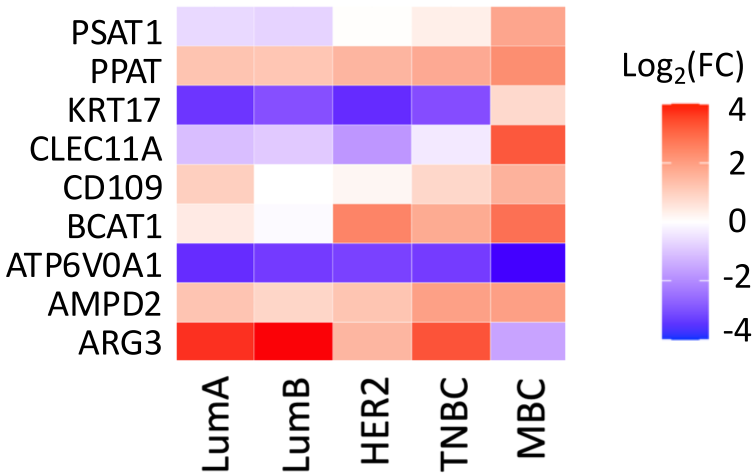


**Supplementary Figure S3. A subset of significantly altered proteins with conserved fold changes in all breast cancer subtypes.** A) Heatmap of the log2(FC) for the 576 significantly altered proteins with conserved differences in all breast cancer subtype comparisons with disease-free tissue. B) A heatmap of the significantly altered protein groups with fold changes that are correlated with breast cancer subtype is displayed. C) A heatmap of 'stemness' proteins uniquely altered in MBC but not in other subtypes. All proteins shown in (A) & (B) are significantly altered with a q-value ≤ 0.001.
