## Supplementary material for "Proteomic Analysis of Breast Cancer Subtypes Identifies Stromal Protein Profiles that Contribute to Aggressive Malignant Behavior": Suppl Figure 4

### Supplementary Figure S4

A **Proteins with the GO-term ”extracellular”**

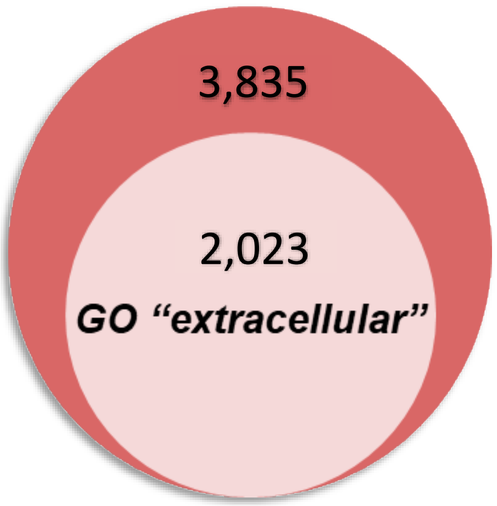


B **MatrisomeDB Protein Distribution**

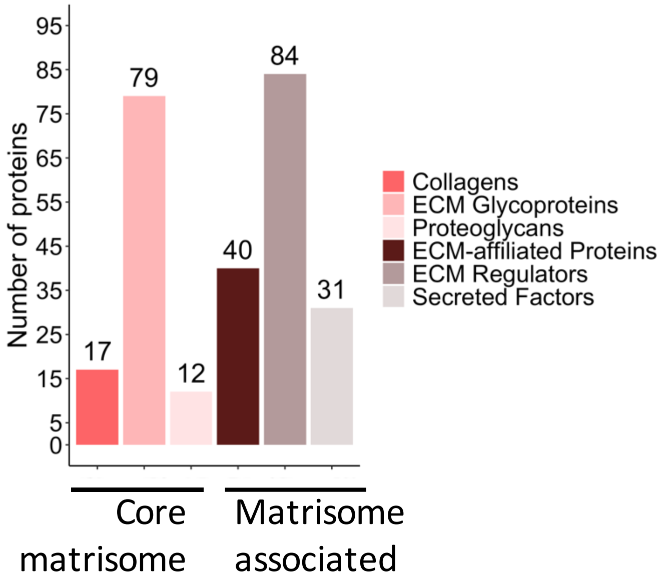


C **MatrisomeDB Common Significantly Altered Protein Distribution**

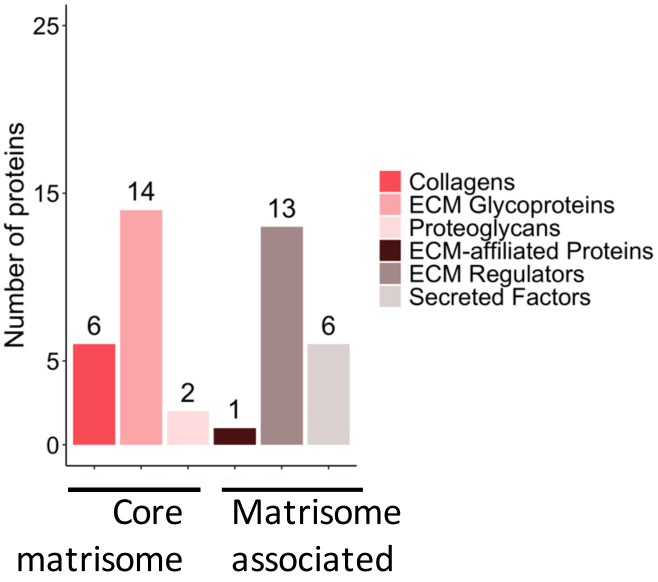


**Supplementary Figure S4. Extracellular protein groups are quantified from FFPE specimens.** A total of 2,023 protein groups from all 5,585 protein groups quantified with 2 or more unique peptides have the GO-term ”extracellular”. The number of quantified protein groups contained in each ECM category from the human MatrisomeDB are shown in a histogram for B) all extracellular proteins identified and C) the subset of 42 extracellular matrix proteins that are significantly altered in all breast cancer subtypes when compared to disease free tissue.
