## Supplementary material for "Proteomic Analysis of Breast Cancer Subtypes Identifies Stromal Protein Profiles that Contribute to Aggressive Malignant Behavior": Suppl Figure 5

### Supplementary Figure S5

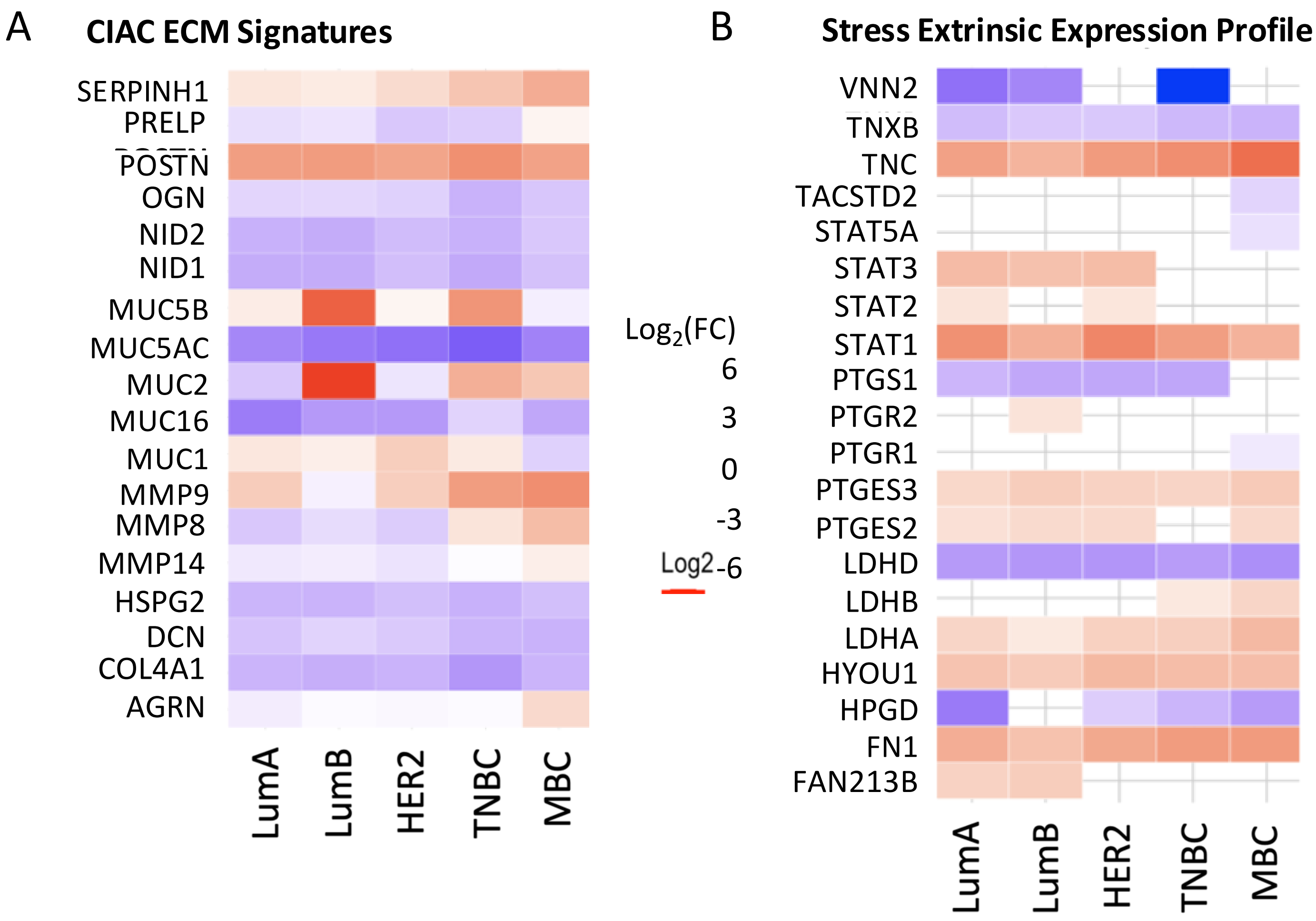


**Supplementary Figure S5. Epithelial stress and chronic inflammation-associated cancer signatures are significantly altered in breast cancers.** A) Heatmap demonstrating regulation of proteins in the epithelial stress extrinsic expression profile in breast cancer.[11] B) Heatmap showing regulation of chronic inflammation-associated cancer extracellular matrix (ECM) signatures in breast cancer subtypes.
