## Supplementary material for "Proteomic Analysis of Breast Cancer Subtypes Identifies Stromal Protein Profiles that Contribute to Aggressive Malignant Behavior": Suppl Figure 6

### Supplemental Figure S6

A **WGCNA Cluster Dendrogram**

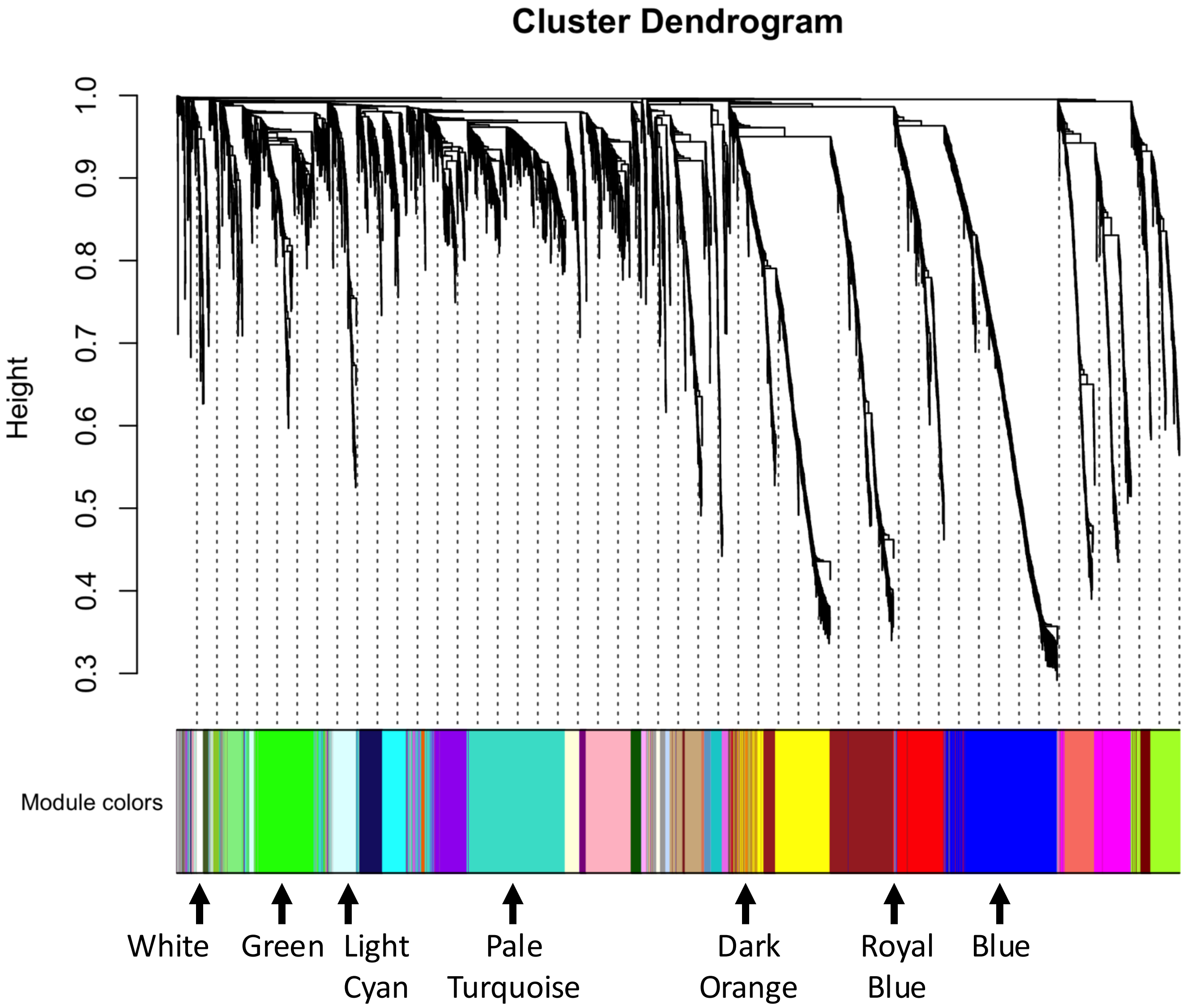


**Supplementary Figure S6. Hierarchical Clustering of Gene Expression Profiles in Breast Cancer FFPE Samples Using Weighted Gene Co-expression Network Analysis (WGCNA).** The dendrogram in (A) illustrates the hierarchical clustering of gene expression profiles derived from formalin-fixed paraffin-embedded (FFPE) breast cancer samples. Utilizing Weighted Gene Co-expression Network Analysis (WGCNA), distinct protein clusters were identified and grouped into modules based on variations in gene expression patterns that correspond to different molecular subtypes of breast cancer. Arrows pointing to module colors of interest highlight, from left to right, the white, green, light cyan, pale turquoise, dark orange, royal blue, and blue modules which are correlated with breast cancer subtypes or disease-free tissue.
