## Supplementary material for "Proteomic Analysis of Breast Cancer Subtypes Identifies Stromal Protein Profiles that Contribute to Aggressive Malignant Behavior": Suppl Figure 7

### Supplementary Figure S7

A **MBC vs. Disease-Free Unique Up-Regulated Biological Processes**

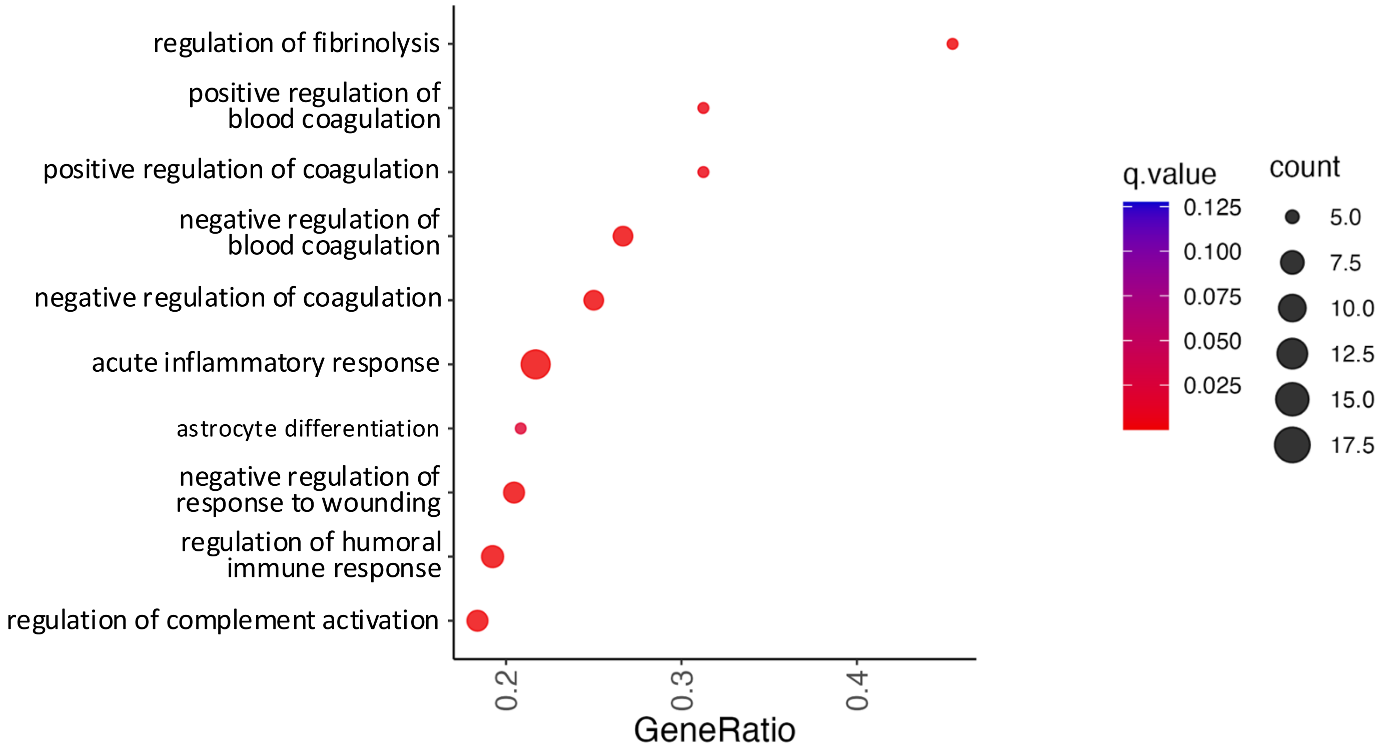


B **MBC vs. Disease-Free Unique Up-Regulated Molecular Functions**

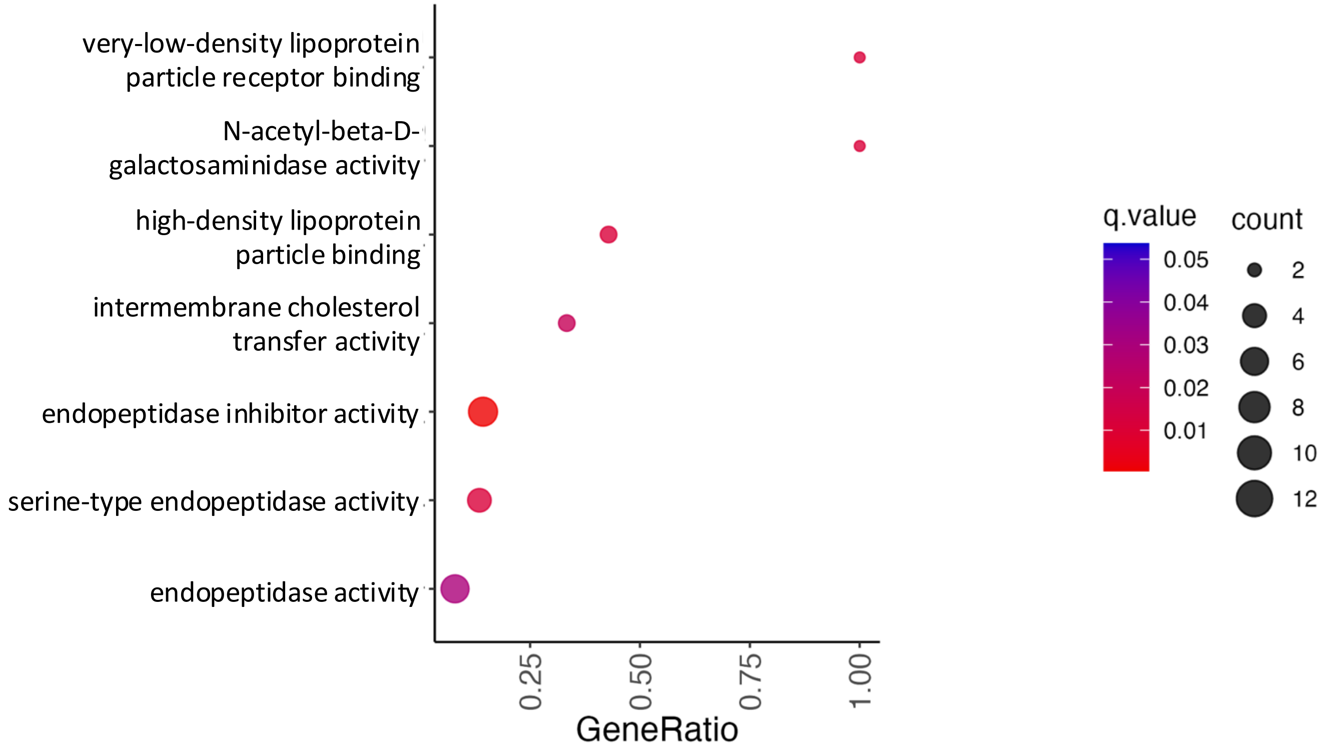

### Supplemental Figure S7

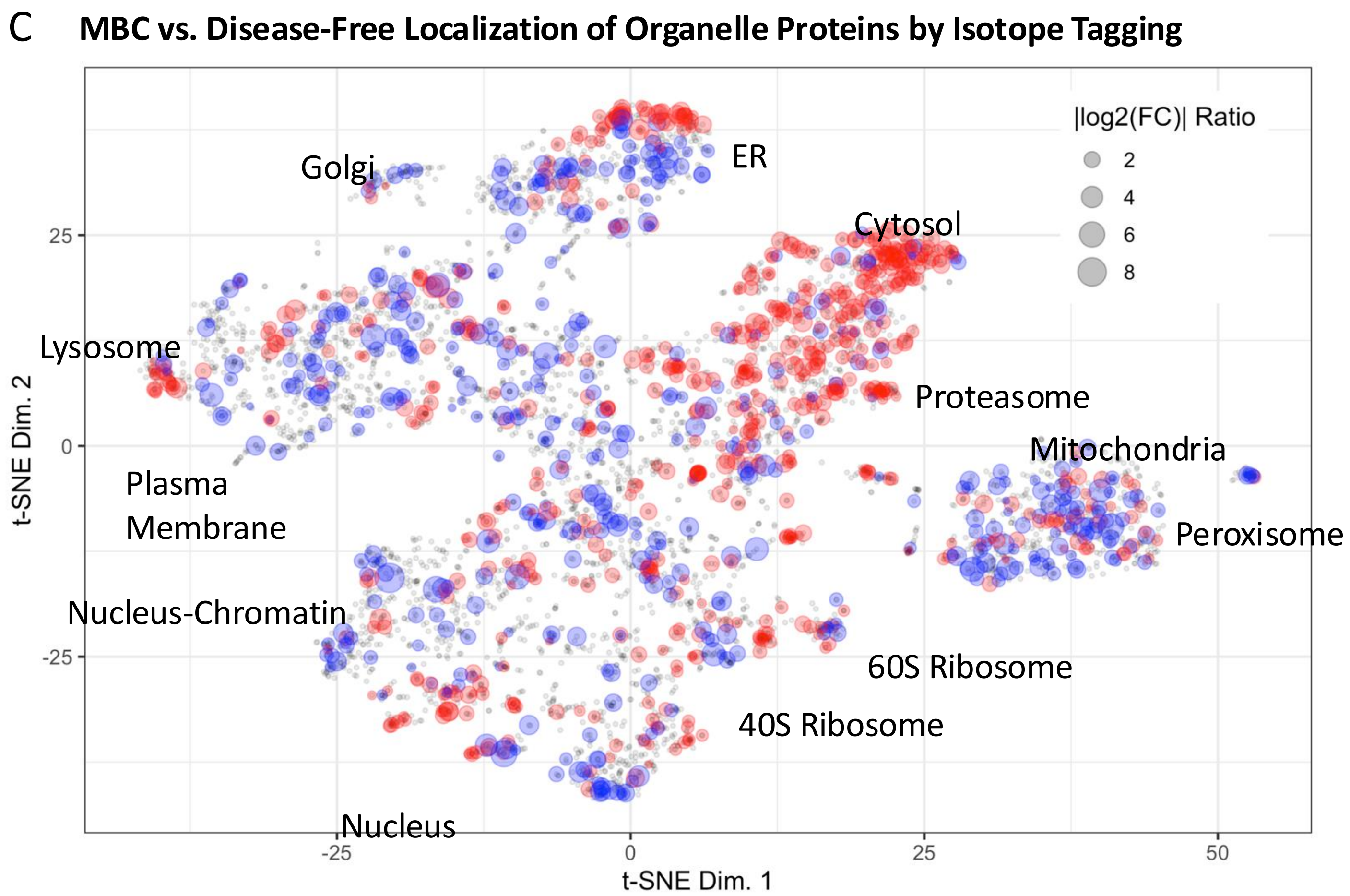


**Supplementary Figure S7. Significantly altered proteins unique in the MBC comparison with disease-free tissue demonstrates disruption of the ECM that is common in chronic inflammation associated cancers.** A) A dotplot shows KEGG biological processes up-regulated uniquely in MBC vs. disease-free tissue. B) Dotplot of the KEGG molecular functions uniquely up-regulated in MBC vs. disease-free tissue. C) Significantly altered protein groups in the MBC vs. disease-free comparison are overlayed onto the Localization of Organellar Proteins by Isotope Tagging plot for human U2OS cells to demonstrate organelle-wide proteome changes in MBC.[23, 24]
