## Supplementary material for "Proteomic Analysis of Breast Cancer Subtypes Identifies Stromal Protein Profiles that Contribute to Aggressive Malignant Behavior": suppl methods

**Supplemental Methods**

*FFPE Breast Tissue Multiplex Immunohistochemistry (mIHC)*

Slides with five-micron thick breast tissue sections were baked at 60^o^C overnight, deparaffinized in two baths of xylene(s) (10 min each) and rehydrated in graded ethanol: 100%, 100%, 95%, 85%, 70% (5 min each), and finally in distilled water. Endogenous peroxidase and autofluorescence quenching was achieved by incubating slides in PBS containing 4.5% H_2_O_2_ and 10 mM NaOH under a bright light for 45 min. Heat-induced antigen retrieval was carried out in citrate buffer, pH 6.0 (Sigma-Aldrich) at 95^o^C for 10 min. Non-specific antibody binding was blocked using Background Sniper buffer (Biocare Medical). Tissue sections were incubated for 1 hour at room temperature (RT) with a first primary antibody (anti-SERPINH1/HSP47, Abcam, rabbit monoclonal antibody clone EPR4217, cat#ab109117, dilution 1/2000; or anti-MPO, Abcam, rabbit monoclonal antibody clone SP72, cat#ab93665, dilution 1/4000; or anti-ECP (RNase3), Cell Signaling Technology, rabbit monoclonal antibody clone E6U5M, cat#24357S, dilution 1/5000) in antibody diluent reagent (ThermoFisher, cat#50-750-38) then for 30 min in pre-diluted MACH 2 Horseradish peroxidase (HRP) labeled goat anti-rabbit micro-polymer secondary antibody (Biocare Medical, cat# RHRP520). The slides were washed in TNT buffer (0.1 M Tris-HCl pH 7.5, 0.15M NaCl, and 0.05% Tween-20), and the signal was developed using Tyramide Signal Amplification (TSA) Plus Fluorescein solution (Akoya Biosciences, cat# NEL741001KT) for 2 min. Removal of the primary antibody complex was achieved by heating in 95 ^o^C citrate buffer pH 6.0 for 5 min. Tissue sections were blocked again with Background Sniper, incubated overnight at 4 ^o^C with a second set of primary antibodies (anti-vimentin, Cell Signaling Technology, rabbit monoclonal antibody clone D21H3, cat#5741, dilution 1/2500; or anti-ITGAM/CD11b, Cell Signaling Technology, rabbit monoclonal antibody clone D6X1N, cat#49420, dilution 1/2000) in antibody diluent reagent then for 30 min in pre-diluted MACH 2 conjugated anti-rabbit secondary antibody. After washes in TNT buffer, the signal was developed using TSA Plus Cyanine 3 (Cy3) solution (Akoya Biosciences, cat# NEL744001KT) for 3 min. Removal of the second primary antibody complex, blocking and incubation for 1h at RT with CD45 antibody (Cell Signaling Technology, rabbit monoclonal antibody clone D9M8I, cat#13917, dilution 1/2000) for the MPO module were performed as described above. The signal was developed using TSA Plus Cyanine 5 (Cy5) solution (Akoya, cat#NEL745001KT) for 12 min. After washes in TNT buffer, nuclei were counterstained with 3 µM DAPI in PBS for 5 min, washed in distilled water, and mounted with Vectashield HardSet Mounting Medium (Vector Laboratories, cat#H-1400). Slides stained with the 3 mIHC marker modules described above were imaged using a BZ-X800 fluorescence microscope (Keyence). Each multiplex-stained specimen was first scanned at low resolution using 4x magnification, and then imaged at high resolution at a 20x magnification. Unless specimens were too small, at least ten independent 20x fields, randomly distributed throughout the tumor region to account for intratumor heterogeneity, were imaged. For sections of disease-free breast tissue, ductal and lobule containing fields were randomly selected in the tissue section.  ImageJ (version 2.14.0/1.54f) was used for image preparation, and CellSeg (1), a Convolutional Neural Network trained model for nuclear segmentation, was used to facilitate nuclear mask generation and staining intensity quantification in nuclear and perinuclear areas. Parameters for CellSeg were set as follows: overlap = 80, threshold = 15, boost = auto, increase factor = 2, growth pixels = 2, growth method = 'Sequential', and should compensate = True. Mean pixel value for each protein of interest was calculated for downstream analysis. Pixel intensity thresholds to identify “positive” cells were as follows: 'SERPINH1': 36, 'Vimentin': 20, 'MPO': 32, 'RNAse3': 32, and 'ITGAM': 25. Pairwise comparisons of number of double-positive cells (SERPINH1/VIM+, MPO+/ITGAM+ or RNAse3+/ITGAM+) per mm2 of tissue area between each breast cancer subtype and disease-free breast were carried out using Prism (version 10.2.0). Two-tailed p-values were calculated using Mann-Whitney tests.

*Further Data Processing and Visualizations*

For biological pathway analysis the Database for Annotation, Visualization, and Integrated Discovery (DAVID, accessed 12/02/2022) was utilized for over-representation analysis (ORA) of significantly altered quantifiable proteins (|log2(FC)| ≥ 0.58 & q-value < 0.001) to determine which KEGG pathway gene ontology terms were enriched in these samples (2). The ggplot2 package in R (version 4.0.5; RStudio, version 1.4.1106) was used to visualize significantly enriched biological processes from each comparison (3). Localization of Organellar Proteins by Isotope Tagging (LOPIT) maps were generated with the pRolocData and pRoloc Bioconductor packages in R (4, 5). The LOPIT map was created using the t-SNE machine learning algorithm to reduce the multi-dimensional human U2OS cell dataset and cluster proteins by similarities, overlaying the points with fold change data comparing breast cancer subtypes to disease-free tissue (4-6). Weighted gene correlation network analysis was performed using the WGCNA package in R (7). The power was set to 5 with a minimum module size of 7 proteins per module. Modules of interest were selected based on the correlation to the breast cancer subtype.

*Experimental Design and Statistical Rationale*

In this study, we used patient-derived FFPE tissues (n = 42) consisting of 7x luminal A, 7x luminal B, 7x Her2+, 7x TN and 7x MBC cancer tissues, as well as 7 disease-free RM specimen used as reference. Multiplex immunohistochemistry experiments were conducted with one technical replicate per biological replicate. Stained slides were imaged using a BZ-X800 fluorescence microscope (Keyence). As described above, unless specimens were too small, at least ten independent, randomly distributed 20x fields were imaged. ImageJ (version 2.14.0/1.54f) was used for image preparation, and CellSeg (1), a Convolutional Neural Network trained model for nuclear segmentation, was used to facilitate nuclear mask generation and staining intensity quantification in nuclear and perinuclear areas. Mean pixel value for each protein of interest was calculated for downstream analysis. Pixel intensity thresholds to identify “positive” cells were as follows: 'SERPINH1': 36, 'Vimentin': 20, 'MPO': 32, 'RNAse3': 32, and 'ITGAM': 25. Pairwise comparisons of number of double-positive cells (SERPINH1/VIM+, MPO+/ITGAM+ or RNAse3+/ITGAM+) per mm^2^ of tissue area between each breast cancer subtype and disease-free breast were carried out using Prism (version 10.2.0). Two-tailed p-values were calculated using Mann-Whitney tests.

Proteomic experiments were conducted with two technical replicate acquisitions per biological replicate. Indexed retention time peptide standards (Biognosys) were spiked into the samples before LC-MS/MS analysis in DIA mode on a ZenoTOF 7600 mass spectrometer. One DIA cycle (2.5 s) was composed of the acquisition of one MS1 scan, followed by the acquisition of 80 variable windows (5-25 m/z) covering the full mass range (m/z 399.5-1000.5) with an overlap of 1 m/z. DIA data were processed in Spectronaut v16 using a peptide-centric approach and a panhuman library containing 149,066 precursors and 10,316 human proteins to retrieve MS2 XIC-based quantification information, (8) as described above, and significantly altered protein groups in each comparison versus disease-free RMs were obtained using a paired t test followed by p-value correction for multiple testing using the Storey method (9).
